# A cryptic hybrid zone reveals the genomic basis of flower colour variation in a plant with a large and complex genome

**DOI:** 10.64898/2026.01.30.700338

**Authors:** Pascaline Salvado, Anaïs Gibert, Slimane Chaib, Delphine Raviglione, Cédric Bertrand, Moaine El Baidouri, Lee Mariault, Christel Llauro, Charlotte Cravero, Isabelle Dufau, William Marande, Marie-Christine Carpentier, Maria Martin, Céline Quélennec, Pascal Gaultier, Jean-Marc Lewin, Pere Aymerich Boixader, Valérie Delorme-Hinoux, Joris A. M. Bertrand

## Abstract

Flower colour is known to be a classic magic trait, undergoing divergent selection pressures exerted by distinct pollinators and involved in reproductive isolation. However, variation at this trait may also results from other eco-evolutionary factors whose interpretation requires throughout context-specific analyses. Here, we investigated the eco-evolutionary causes of pink- and yellow-flowered morphs in *Pedicularis comosa* where their parapatric ranges form a contact zone in the East of the Pyrenees Mountains. First, we generated a near-chromosome-scale reference genome assembly for this species. We then used a Genotyping By Sequencing approach to infer patterns of genetic diversity and differentiation between populations and morphs at a total of 158 individuals from 11 localities. We found that neutral genetic structure is primarily consistent with geography rather than with colour. However, admixture analyses and clines suggest the existence of a cryptic hybrid zone between morphs. Outlier detection methods and examination of locus-by-locus cline features, then allowed to pinpoint candidate *loci* to explain colour variation. To gain insight into the functional aspects of these *loci*, we finally analysed floral transcriptomes and quantified pigments using Liquid Chromatography coupled with Mass Spectrometry (LC-MS) and confirmed the involvement of key genes in the anthocyanin metabolic pathway (e.g. DFR, FLS) and associated pigments (e.g. cyanidin and delphinidin). Our results show that implementing a highly integrative multi-omic approach can allow unraveling the genetic basis and the eco-evolutionary significance of adaptive traits with even very limited previous knowledge, on species with large and complex genomes.

## Introduction

A striking aspect of the Angiosperm radiation is the remarkable diversity of flower colour exhibited by these organisms (Muchhala et al. 2014). The colour patterns displayed by flowering plants vary across the ultraviolet (UV) and visible range (Chittka et al. 1994), thus encompassing the entire colour spectrum capable of being perceived by both humans and other animal pollinators (Menzel and Shmida 1993). The colouration of flowers is generally determined by pigments that are biosynthesised via multiple pathways (Tanaka et al. 2008), but is also subject to variation depending on cell tissue structure and the manner in which light is reflected (Vignolini et al. 2012).

The colour of flowers is an example of a highly labile evolutionary trait, which can differ between sister species (Wesselingh and Arnold 2000; Martén-Rodríguez et al. 2010), among populations of the same species (Streisfeld and Kohn 2005; Cooley et al. 2011), or even among individuals across different life stages. For instance, *Pulmonaria collina*’s flowers undergo a chromatic shift from pink to blue with age, thereby directing pollinators to young and highly rewarding pink flowers (Oberrath and Böhning-Gaese 1999). The predominance of a particular floral colour is generally indicative of adaptation to pollinator-induced selection pressures (Fenster et al. 2004). Indeed, pollinators have long been considered as primary drivers of plant diversification and floral evolution (Dodd et al. 1999; Fenster et al. 2004). This is evidenced by the significantly higher species richness exhibited by animal-pollinated lineages in comparison to abiotically-pollinated lineages (Dodd et al. 1999; Strauss and Whittall 2006).

Colour is one of the most significant cues employed by pollinators in the location, recognition, and discrimination of different flowers (Menzel and Shmida 1993; Schiestl and Johnson 2013). However, mounting evidence suggests that biotic and abiotic non-pollinator agents also play a crucial role in flower colour as a response to selection (Schemske and Bierzychudek 2001). Firstly, it has been demonstrated that temperature and drought stresses tend to favour floral tissues containing UV-absorbing pigments (Rausher 2008), such as anthocyanins. For example, in various species, pink or purple-flowered individuals have been shown to be selected for over white-flowered ones, as anthocyanins have been demonstrated to enhance stress tolerance in vegetative tissues (Strauss and Whittall 2006). Furthermore, studies have shown that anthocyanin-containing morphs induce a reduction in herbivore performance in comparison to anthocyanin-less morphs (Strauss et al. 2004). It has been also demonstrated that light intensity exerts a positive selection pressure on UV floral pigmentation, given its capacity to influence the environment surrounding pollen and ovules. It was posited that larger areas of UV-absorbing pigmentation might serve a protective function by safeguarding pollen from UV-induced damage (Koski and Ashman 2016). Additionally, soil composition has been shown to influence flower colour variation (Schemske and Bierzychudek 2007). Experiments with *Hydrangea macrophylla* demonstrated that pH variations in the soil were able to induce a shift in flower colour from blue to pink (Wallace and Wieland 1980).

The study of flower colour variation in natural populations poses a significant challenge to the field of evolutionary biology, while simultaneously offering a unique opportunity for research. It is evident that such variation can be shaped by a complex interplay of ecological, genetic and historical factors. The interpretation of its eco-evolutionary significance is thus context-specific. Conversely, the investigation of biological systems exhibiting intra-specific flower colour variation at the spatial scale of a local landscape may facilitate the comprehension of microevolution. This phenomenon is especially evident in systems where parapatric morphs form zones of contact, occasionally exhibiting intermediate floral phenotypes in areas of overlap. This observation suggests the possibility of hybridisation. The study of such contact – or hybrid – zones is challenging, as it necessitates the disentanglement of patterns of gene flow, selection, and historical divergence. However, these regions also possess a remarkable informational value, as they constitute natural experiments in which evolutionary processes unfold in real time. Hybrid zones are defined as regions where genetically distinct populations meet and interbreed (Barton and Hewitt 1985), and have been described as ‘natural laboratories for evolutionary studies’ (Hewitt 1988) and ‘windows on evolutionary processes’ (Harrison 1990). These species offer unique opportunities to identify traits implicated in adaptation and reproductive isolation, and to establish links between these traits and their genomic underpinnings (Chase et al. 2017). The genesis of such zones may be either primary, resulting from intergradation along environmental gradients (Endler 1977; Caisse and Antonovics 1978), or secondary, arising from secondary contact following allopatric divergence (Mayr 1942).

For example, research conducted on a monkeyflower species (*Mimulus aurantiacus*) hybrid zone has indicated that pollinator-mediated reproductive isolation constitutes a primary barrier to gene flow between two divergent red- and yellow-flowered ecotypes pollinated by a hummingbird and a hawkmoth respectively (Chase et al. 2017; Stankowski et al. 2022). Another well-known example is the two morphs of snapdragons (*Antirrhinum majus*): the magenta-flowered *A. majus* subsp. *majus* var. *pseudomajus* and the yellow-flowered *A. majus* subsp. *majus* var. *striatum* which come into contact and very locally hybridise in the east of the Pyrenees mountains (Whibley et al. 2006; Field et al. 2025; Pal et al. 2025). In this example, the flower colour differentiation cannot be clearly associated with pollinator-mediated selection pressures, since both ecotypes share common and generalist insect pollinator species (Khimoun et al. 2011). However, it has been demonstrated that specific combinations of climatic and other abiotic-related factors such as elevation could influence the presence of one subspecies instead of the other (Khimoun et al. 2013; Marin et al. 2020). Furthermore, genetic variation between the two subspecies has been found to be predominantly influenced by landscape and the evolutionary history of the species (Pujol et al. 2017).

The present study, focuses on a system that intriguingly reminds *Antirrhinum majus*, involving, in the same geographic context, a contact zone between pink- and yellow-flowered populations but in a phylogenetically distant plant family (Orobanchaceae). The species *Pedicularis comosa* comprises two distinct morphs described as ‘subspecies’: the yellow-flowered *Pedicularis comosa* subsp. *comosa*, widespread across mountain regions of the Palearctic and the pink-flowered *Pedicularis comosa* subsp. *asparagoides*, endemic to the Eastern Pyrenees. The geographic distributions of these two ‘subspecies’ meet but exhibit no apparent overlap, with the potential for a cryptic hybrid zone between the two different morphs occurring in the eastern part of the Pyrenees (Figure 1).

**Figure 1.**
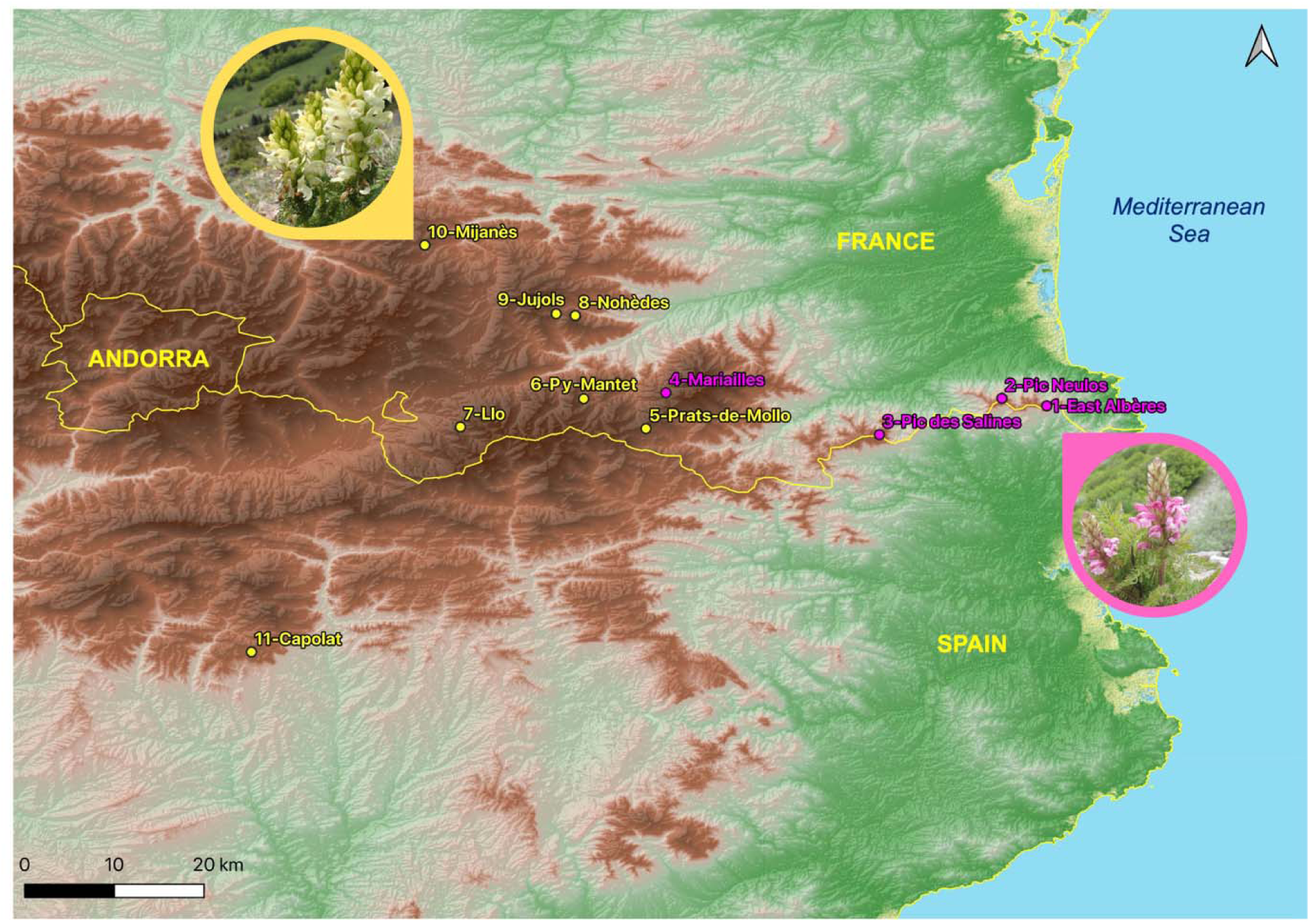
Map representing the geographical location of the 11 localities of the parapatric morphs of *Pedicularis comosa*: *P. c. comosa* (yellow-flowered) and *P. c. asparagoides* (pink-flowered) sampled along a longitudinal transect in the Eastern Pyrenees in this study.

This species is an herbaceous, perennial, allogamous and hemiparasitic plant that reaches 10 to 60 cm in height. Its longevity is not well documented, but it could live for a minimum of 10 years. The flowering period takes place between May and July depending on elevation, and cross-pollination is carried out by bumblebees (*Bombus sp*.). However, the biology and ecology of this representative of the Orobanchaceae family remain to be fully elucidated. Historically, its classification has been subject to variation, with two taxa being either distinguished as distinct *species* or as two subspecies based on floral pigmentation and subtle morphological differences such as the maximum width of foliar segments (narrower in the pink-flowered *P. c.* subsp. *asparagoides*). The latter morph is also endemic to the easternmost part of the Pyrenees, and Species Distribution Modeling (SDM) analyses have shown that its suitable habitat may disappear in the XXI^st^ century because of climate change (Salvado et al., *in press*).

Here we aim to investigate the genetic basis of flower colour variation in *Pedicularis comosa*, to address the eco-evolutionary significance of this trait and to clarify the systematics and the taxonomy of this species. To do so, we first sequenced, assembled and annotated a high-quality reference genome for *P. comosa*. Then, based on ca. 150 individuals sampled in the field at 11 localities (4 pink-flowered ones and 7 yellow-flowered ones), we employed a population genomics approach to i) investigate patterns of neutral genetic variation among populations, ii) look for genomic outlier regions possibly involved in local adaptation and/or reproductive isolation and iii) compare patterns of phenotypic and genetic variation along a longitudinal gradient and their putative clinal nature. To further gain insights into the functional aspects associated with floral colour variation, we finally generated and compared transcriptomic data comprising pink- and yellow-flowered individuals and quantified pigment variation through Liquid Chromatography-Mass Spectrometry (LC-MS) approaches.

## Results

### Genome assembly, annotation, genomic synteny, repeat and transposable elements content

We generated a high-quality, near chromosome-level reference genome for *Pedicularis comosa asparagoides* using a of PacBio Hifi (Supplementary Table S1) and optical maps. We obtained a 2.8 Gb genome composed by 48 scaffolds with an N50 of 161 Mb. The predicted number of protein-coding genes was 48,030 (Supplementary Table S2) in addition to repeats (Supplementary Table S3) and noncoding RNAs (Supplementary Table S2). Approximately 98.4% of the 425 benchmarking universal single-copy (BUSCO) genes were recovered in full-length (Supplementary Table S1). Overall, 71.9% of the *P. comosa* genome is composed of TEs (Supplementary Table S3). As in other angiosperms, Class I elements represent the vast majority of TEs in *P. comosa*, accounting for more than 81%, while Class II elements account for 18%. Among Class I elements, LTR retrotransposons are the most abundant superfamily, representing 76% of total TEs.

### Population genomic dataset

We obtained a total of 602 million read pairs across the 156 individuals (1 721 895 - 9 038 454 of raw reads per ind., mean 3 809 397, SD=1 138 502). We assembled a total of 3,279,982 *loci*. Average read depth per individual ranged from 22.7x to 94.2x based on the combination of parameters we used, described in the Methods. After filtering the data with the *populations* script (from *STACKS*), we finally obtained 14,749 *loci* including 28,729 SNPs.

### Genetic diversity and structuration

All sampled populations significantly deviated from panmixia (mean *G*_IS_ = 0.193, *p* < 0.05), with similar levels of allelic diversity across sites (Table 1). Private allele counts reflected the degree of geographic isolation (Table 1). Multivariate, phylogenetic, and clustering approaches (PCA, Maximum-Likelihood tree reconstruction, sNMF, ADMIXTURE; Figure 2, Supplementary Figures S2–S3) consistently recovered three geographically coherent groups: Albères (populations 1–3), Canigou (4–7), and Madres (8–10). Capolat (11) was highly divergent from all others localities, with pairwise *G*_ST_ values > 0.345 (Supplementary Table S5). A significant isolation-by-distance pattern (Mantel *r* = 0.517, *p* = 0.015; Supplementary Figure S4) suggests that gene flow is generally limited between massifs. Pic des Salines (3) consistently appeared intermediate between the Albères (1-3) and Canigou (4-7) massifs in the PCA, phylogenetic tree, FineRADstructure, and sNMF analyses. At *K* = 4 with sNMF, this population showed nearly equal ancestry from Albères (light blue) and Canigou (pink) clusters, while ADMIXTURE indicated a slightly different pattern, placing 4-Mariailles as the more admixed population at *K* = 3 (Figure 2, Supplementary Figure S2). Despite these differences, both methods highlight localised admixture between Albères and Canigou massifs. These patterns collectively reveal strong spatial structuring, with genetic connectivity largely restricted to contact zones.

**Figure 2.**
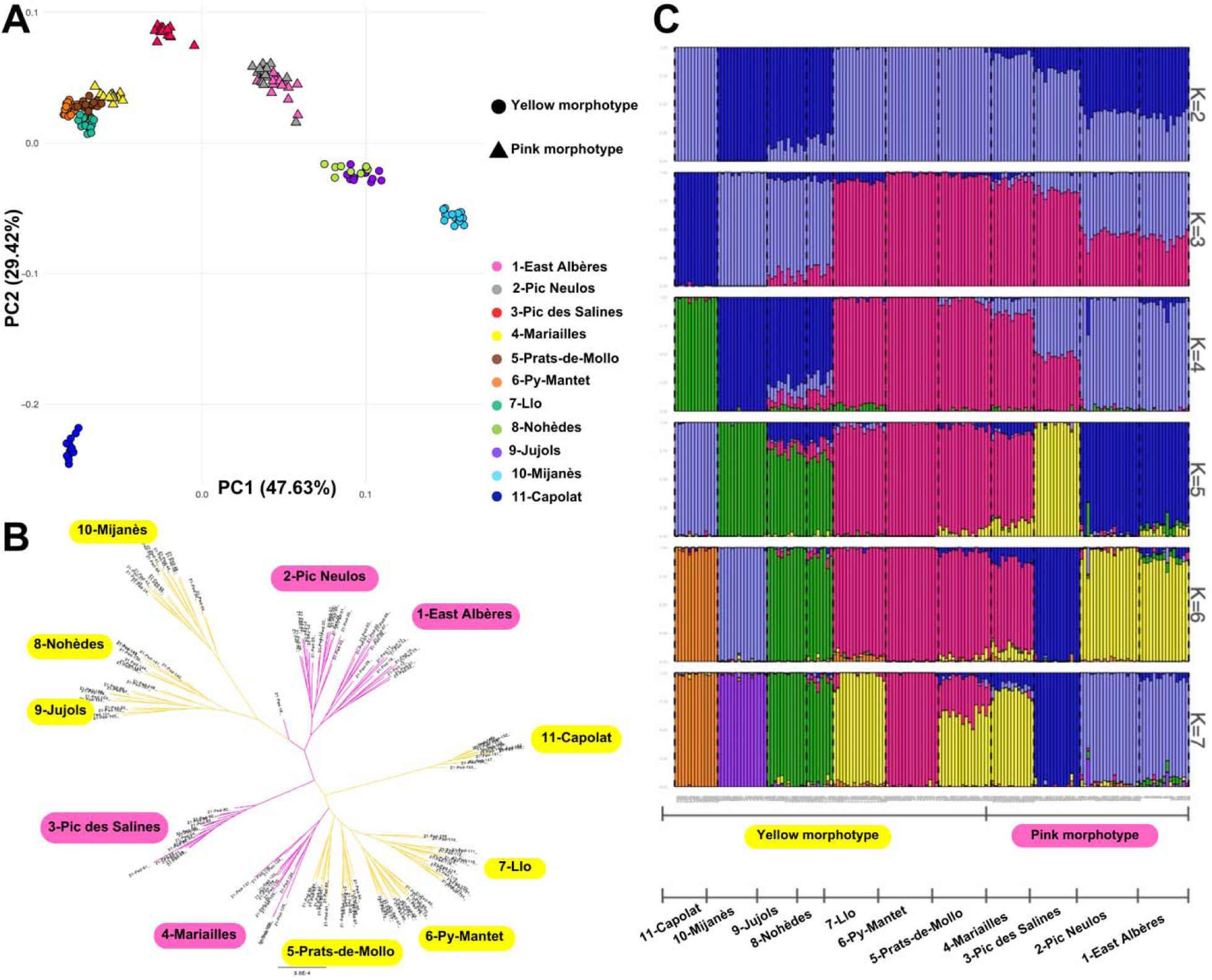
Phylogeny, population structure and admixture. **A)** Principal Component Analysis (PCA) biplot of the two first axes (PC1) and (PC2) summarizing 47.63% and 29.42% of the total genetic variance, respectively. This PCA was computed based on a matrix of 156 individuals (as rows) and 28,729 SNPs (as columns) **B)** Unrooted phylogenetic tree of 156 individuals on the basis of alignments of 14,749 nGBS *loci* and 28,729 SNPs **C)** Barplots displaying individual ancestry coefficients obtained from sNMF for 156 individuals and 28,271 for *K*=2 to *K*=7. The optimal value of *K* was found to be 6.

**Table 1.** Geographic coordinates (in °) and elevation (in m above sea level) of the sampling localities, sample size, observed (*H*_O_) and expected (*H*_E_) heterozygosity, deviation from panmixia *G*_IS_, total number of alleles (*A*), and number of private alleles (*A*_p_) per locality

| Locality | Latitude | Longitude | Elevation | $N$ | $H_o$ | $H_E^a$ | $G_{IS}$ | $A$ | $A_p$ |
| --- | --- | --- | --- | --- | --- | --- | --- | --- | --- |
| 1-East Albères | 42.472 | 3.017 | 1011 | 15 | 0.180 | 0.231 | 0.218 | 18620 | 121 |
| 2-Pic Neulos | 42.484 | 2.947 | 1205 | 18 | 0.165 | 0.203 | 0.187 | 16000 | 131 |
| 3-Pic des Salines | 42.426 | 2.751 | 1359 | 14 | 0.127 | 0.149 | 0.149 | 11532 | 140 |
| 4-Mariailles | 42.493 | 2.41 | 1802 | 13 | 0.128 | 0.184 | 0.303 | 14607 | 95 |
| 5-Prats-de-Mollo | 42.436 | 2.378 | 2134 | 16 | 0.144 | 0.187 | 0.231 | 16819 | 76 |
| 6-Py-Mantet | 42.483 | 2.279 | 2007 | 16 | 0.141 | 0.173 | 0.183 | 14527 | 73 |
| 7-Llo | 42.438 | 2.082 | 1601 | 16 | 0.138 | 0.177 | 0.217 | 14515 | 96 |
| 8-Nohèdes | 42.616 | 2.265 | 1641 | 8 | 0.184 | 0.212 | 0.135 | 12175 | 73 |
| 9-Jujols | 42.619 | 2.235 | 1690 | 12 | 0.190 | 0.232 | 0.182 | 15437 | 96 |
| 10-Mijanès | 42.729 | 2.025 | 1590 | 15 | 0.159 | 0.210 | 0.241 | 15134 | 195 |
| 11-Capolat | 42.078 | 1.748 | 970 | 13 | 0.103 | 0.112 | 0.079 | 8178 | 163 |
|  |  |  |  | 156 | 0.151 | 0.188 | 0.193 |  |  |

Population genetic statistics were computed on the basis of 28,729 SNPs. Statistically significant values are indicated in bold ( $p < 0.05$ ). $H_E$ : unbiased expected heterozygosity measured following Nei (1987), in Genodive.

### Introgression analysis

Triangle plots (Figure 3A) revealed no individuals with high heterozygosity (all < 0.3) and thus, putative F1 hybrids, but individuals from 3-Pic des Salines exhibited hybrid indices near 0.5, consistent with admixture at *K* = 4 (Figure 2C) between the Alberes massif (1–2) and the Canigou (4–7). These patterns confirm localised gene flow, primarily linking Albères (1-2) and Canigou (4-7), with secondary connections to Madres (8-10) found by *D*suite (Supplementary Figure S5).

**Figure 3.**
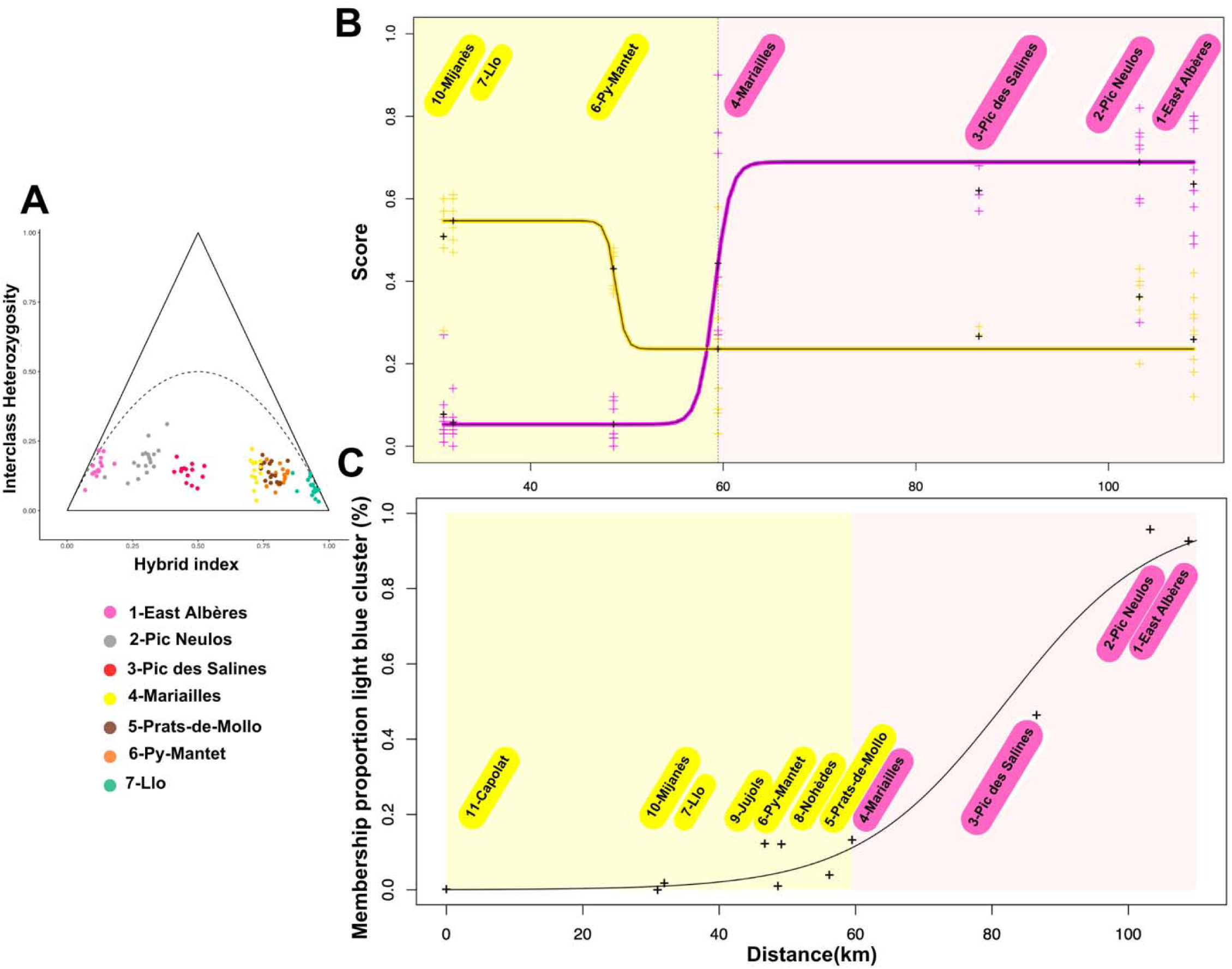
**A)** Triangle plot drawn with triangulaR with the 7 populations constituting the contact zone from the East to the West side of the Pyrenees. **B)** Phenotypic clines fitted with HZAR on the magenta and yellow scores extracted from the calibrated pictures. **C)** Genetic clines fitted with HZAR on the membership proportion of the light blue cluster displayed by SNMF with *K*=4.

### Cline analysis

Phenotypic clines for flower colour (Figure 3B) revealed steep, sigmoid-shaped transitions for both yellow (Y) and magenta (M) scores, centered at 49.55 km and 58.94 km from 11-Capolat, with very narrow widths (7.37 km and 4.41 km; Supplementary Table S6). In contrast, clines fitted to genomic hybrid indices (Q) derived from sNMF and ADMIXTURE (Figure 3C, Supplementary Figure S6) were broader and shifted eastward, with centers near 82 km along the transect (82.11 km and 79.82 km, respectively; Supplementary Table S6). Pic des Salines (3) showed inconsistent placement: central in the sNMF cline but displaced toward the Albères end (1-2) in the ADMIXTURE cline. This mismatch suggests that the two methods differ in sensitivity to genetic structure, with ADMIXTURE capturing a sharper transition between the Canigou (4-7) and Albères (1-2) massifs, while sNMF highlights a more gradual admixture gradient.

### Outlier detection analysis

Geographic cline analyses revealed a mean cline center at around 55 km and a mean width of 29.6 km (Supplementary Figure S7). We identified two strong candidates with steep clines (Figure 4B, Supplementary Table S7): Dihydroflavonol-4-reductase (DFR) and a UDP-glycosyltransferase, both also highlighted by AMOVA as highly differentiated between groups (here, morphs). GWAS additionally identified another UDP-glycosyltransferase (7deoxyloganetic acid glycosyltransferase) associated with magenta pigmentation score via the BLINK model. These candidate *loci* exhibit narrow clines (widths <5 km), centered between 50–60 km (Supplementary Figure S8), overlapping the phenotypic transition zone.

**Figure 4.**
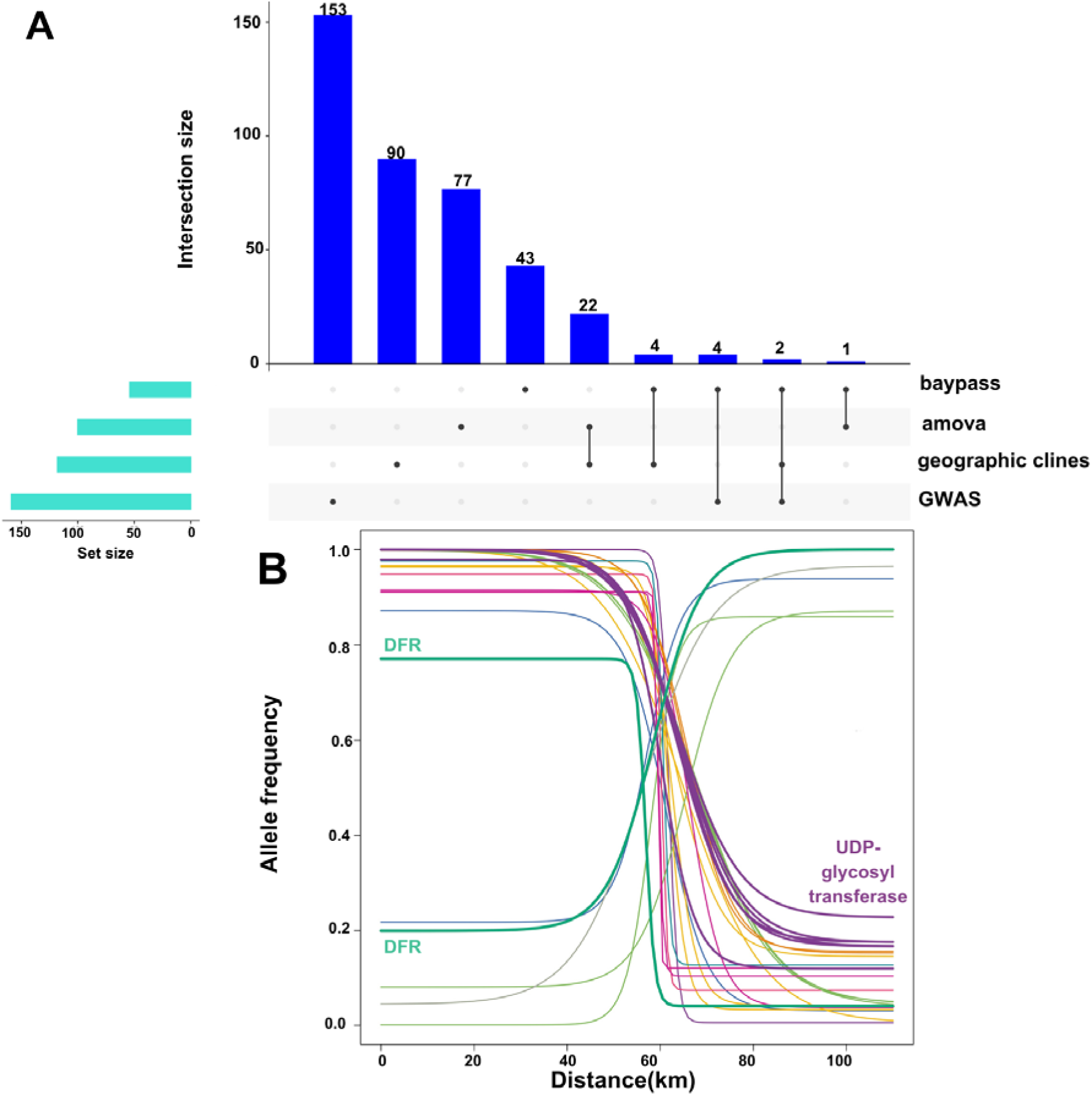
**A)** Upset plot with four different methods to find genomic outliers involved in colouration. **B)** Twenty-eight steepest geographic clines fitted with HZAR. Scaffolds are displayed in different colours. The two clines corresponding to the DFR region are represented in bold and belong to the Scaffold 15,494. The mean of the center and the width of these steepest clines are 62.51 km and 17.68 km respectively.

### Transcriptomes analysis and differential gene expression

The 20 samples of *Pedicularis comosa* sequenced have a mean total number of reads of 24,890,677 M/ind. (18,863,877 min and 41,939,666 max). Mapping rate onto the reference genome was high with a mean of 85,26% (Supplementary Table S8). Among KEGG-annotated metabolic pathways involved in anthocyanin biosynthesis, only DFR and FLS2 were significantly differentially expressed between morphs (Figure 5A). Expression contrasts between pink and yellow flowers were consistent with pigment biosynthetic pathways involving these two genes (Figure 5C).

**Figure 5.**
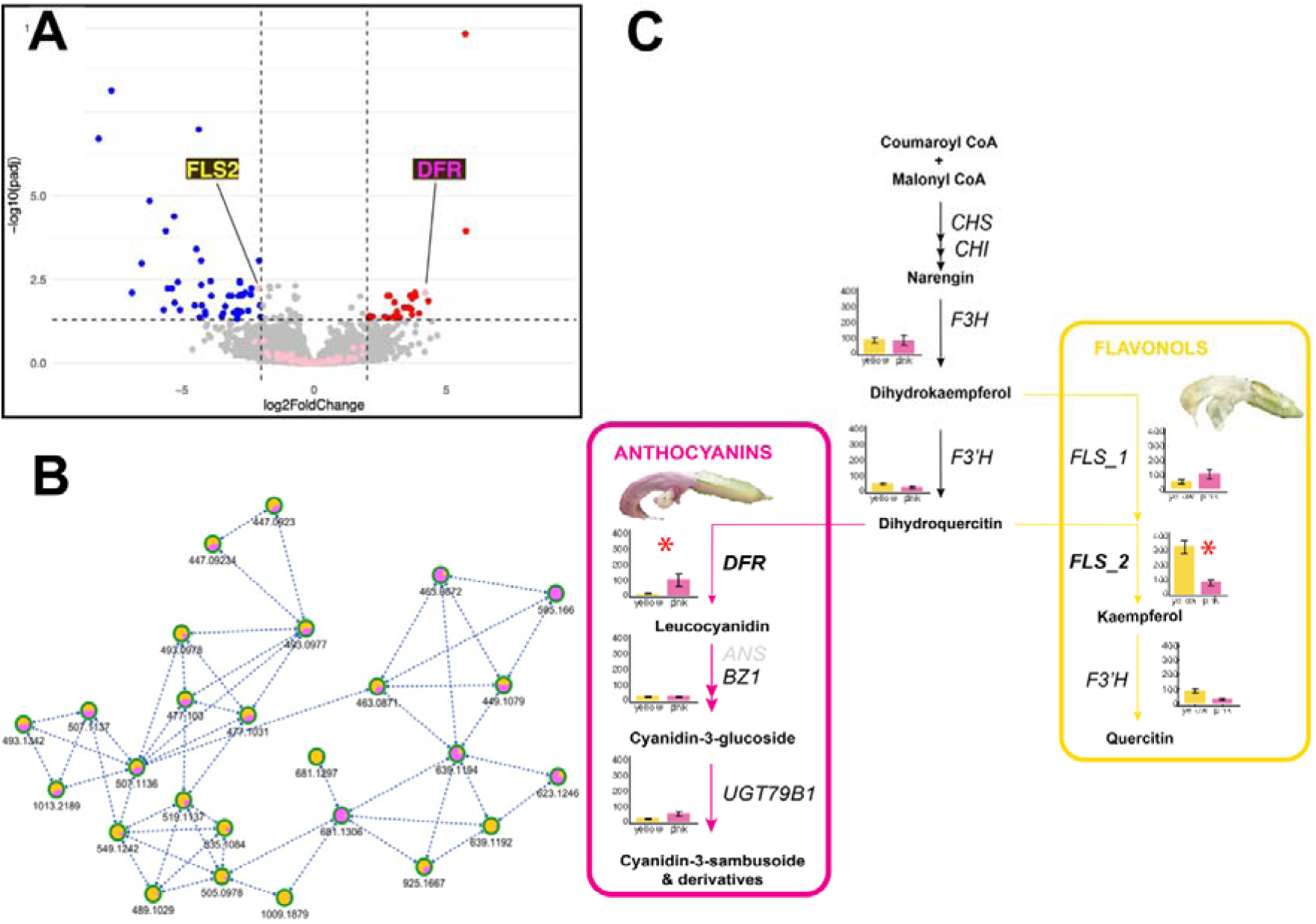
**A)** Volcano plot showing differentially expressed genes between pink- and yellow-flowered individuals of *P. comosa*. Genes upregulated in pink individuals are shown in red, while those upregulated in yellow individuals appear in blue. Known candidate genes involved in pigment biosynthesis, previously identified in other plant systems, are highlighted in light pink. The positions of DFR and FLS1 are indicated. **B)** Molecular network of flavonoid-related metabolites detected by LC–MS in positive ionization mode. Nodes represent molecular features (m/z) and edges indicate spectral similarity. Node coloration reflects relative abundance in pink and yellow floral morphs. The most abundant ion (m/z 595.17), annotated as a cyanidin–coumaroyl–rhamnoside, clusters with closely related cyanidin derivatives and is specific to pink flowers, whereas several flavonoid-related ions enriched in yellow morphs form distinct clusters. **C)** Simplified representation of the anthocyanin and flavonol biosynthesis pathways, including the enzymes involved and their expression levels in the two floral morphs. Gene expression is shown as normalised read counts for each morphotype (pink vs. yellow). Among all the genes shown, only DFR (anthocyanin pathway) and FLS2 (flavonol pathway) were significantly differentially expressed, with higher expression in the pink and yellow morphs, respectively (indicated by asterisks)

### Pigment analysis

LC–MS analyses revealed that pink floral pigmentation in *Pedicularis comosa* is associated with two major anthocyanin-derived compounds, one of which has not previously been reported in the genus. One compound was detected in both floral morphs but was significantly more abundant in pink flowers, whereas the second compound was exclusively detected in pink morphs and entirely absent from yellow flowers (Wilcoxon tests, *p* < 0.05) (Figure 5B).

UV–visible spectra and MS/MS fragmentation patterns indicated that these metabolites correspond to cyanidin- and delphinidin-derived anthocyanins. In particular, the pigment specific to pink morphs was annotated as a cyanidin-based acylated glycoside, while the second compound corresponded to a delphinidin-derived rutinoside. Together, these results indicate that floral colour divergence in *P. comosa* primarily reflects qualitative and quantitative differences in anthocyanin composition, with cyanidin- and delphinidin-derived pigments being strongly enriched in pink-flowered individuals (Figure 5B).

## Discussion

Our study provides key insights into the evolutionary dynamics of *P. comosa*. First, we resolved the taxonomic debate by showing that flower colour does not fully correspond to neutral genetic differentiation, supporting the recognition of a single species with two colour morphs rather than two different taxa. Second, we documented that the contact zone in the Canigou massif, is consistent with a cryptic hybrid zone, marked by both phenotypic and genetic intermediacy. Third, we uncovered candidate genomic regions involved in determining the differences between morphs. Finally we proposed a verbal mechanistic model for flower colour divergence thanks to an integrative approach combining population genomics, geographic and genomic cline analyses, transcriptomics, and metabolomics (pigment analyses). This highlights the power of integrative approaches to rapidly gain insights into the genetic architecture of key traits, even in challenging non-model organisms for which we possess very limited preliminary knowledge.

### Flower colouration has no taxonomic significance in *Pedicularis comosa*

Our results show that the colour of the flowers does not fully reflect the geographic patterns of neutral genetic differentiation among *Pedicularis comosa* populations in the Eastern Pyrenees. Historically, the pink- and yellow-flowered morphs of *P. comosa* L., 1753 have been considered as two distinct taxa. Initially considered as forming a distinct species (*Pedicularis asparagoides* Lapeyr., 1803), the pink-flowered populations, which are endemic to the eastern part of the Pyrenees, have more recently been regarded as a subspecies: *P. comosa* subsp. *asparagoides* (Lapeyr.) P. Fourn., due to the limited number of traits (other than flower colour) allowing it to be distinguished from the yellow-flowered *P. comosa* (also referred to as *P. comosa* subsp. *comosa*). This distinction between the two taxa was further supported by the geographic distribution of yellow- and pink-flowered populations, whose ranges come into contact but do not overlap (Supplementary Figure S9). Notably, there are no populations displaying both yellow- and pink-flowered individuals where they meet. The phylogenetic hypothesis we proposed herein shows that pink- and yellow-flowered populations do not form reciprocally monophyletic clades. Instead, we found that the overall genetic structure was found to be more consistent with geography (*i.e.* mountain massifs) than with flower colour although the two features generally covary in this system. Localities with either yellow-flowered plants (*i.e.* 5-Prats-de-Mollo, 6-Py-Mantet, 7-Llo) or pink-flowered plants (*i.e.* 4-Mariailles) nearby the contact zone of the Canigou-Puigmal massif form a unique clade/genetic cluster. Consequently, our findings provide compelling evidence for a single species comprising two distinct colour morphs that are geographically parapatric yet display a more complex neutral genetic structure that has been primarily shaped by the rugged topography of the landscape, rather than two well unambiguously defined taxa (subspecies or species). This suggests that the eco-evolutionary processes and the genes underlying flower pigmentation are not fully aligned with the overall historical signal.

### A “cryptic” hybrid zone rather than a simple contact zone between the two morphs of *P. comosa*

Our results provide strong evidence that the transition between pink and yellow-flowered morphs of *P. comosa* corresponds to a hybrid zone, rather than a simple contact zone where morphs meet without interbreeding. Phenotypic clines for flower colour traits (yellow and magenta scores, Figure 3A) are remarkably sharp and narrow (less than 10 km wide) and are centered between 50 and 60 km from the westernmost locality (11-Capolat), precisely in the 4-Mariailles/5-Prats-de-Mollo area. Such steep clines (sigmoid S-shaped) are a classical signature of strong divergent selection maintaining phenotypic differences in the face of gene flow. This pattern is further corroborated by field observations, which reveal some individuals at 4-Mariailles with intermediate floral phenotypes (e.g., pale pink, whitish flowers, or pink petals with white patches), suggesting ongoing or at least recent hybridisation. Yet, an examination of individual colour scores at this locality also show a trend towards more extreme values (Figure 3A) for both pink and yellow flowers, an element that is consistent with a phenomenon of potential reinforcement (Barton and Hewitt 1985; Howard et al. 1993; Servedio and Noor 2003), supporting the idea that flower colour is under divergent selection pressures there (and/or that hybrids may be less fit that their parents).

In contrast, the geographic cline found with the membership proportion to clusters with sNMF and ADMIXTURE are wider (33-43 km) and shifted eastward, with centers located around 80 km, near 3-Pic-des-Salines. While this discrepancy might suggest a lack of alignment between phenotype and genomic structure, a closer look reveals a more nuanced picture. At *K*=4, sNMF shows 3-Pic-des-Salines as genetically admixed (*i.e.* 46% for the light blue cluster and 53% for the pink cluster); and at higher *K* values (*K*=5, 6, 7), both sNMF and ADMIXTURE consistently identify it as a distinct genetic cluster. Thus, genetic singularity is further supported by PCA, phylogenetic tree, triangle plot and FineRADstructure (Figure 2A and B, Figure 3C and Supplementary Figure S3 respectively), all of which place this population in an intermediate position between the Albères and Canigou massifs. These patterns may be shaped by geographic isolation (due to the Rome Valley and Tech River) or reflect methodological sensitivity to complex geographic signals (e.g. asymmetric gene flow, ghost introgression of post-admixture drift (Toyama et al. 2020; Pang and Zhang 2024). Moreover, the ecological context may play a key role in shaping this pattern. It has been shown that clines in hybrid zones often become “trapped” at ecological discontinuities, where population densities are naturally low (see Delahaie et al. 2017). In our system, the Tech Valley, a low-elevation zone nowadays unsuitable for *P. comosa*, may act as such a barrier. This could explain why the center of the genomic cline appears displaced relative to the phenotypic cline and suggests that the hybrid zone in *P. comosa* may be geographically unstable, with historical or ongoing shifts in its position.

Crucially, when examining geographic clinal patterns of SNP allele frequencies (Figure 4B, Supplementary Table S7, Supplementary Figure S7 and S8), we find that a substantial proportion of *loci* exhibit sharp and narrow allele frequency transitions centered between 50 and 60 km, again in the 4-Mariailles/5-Prats-de-Mollo regions, where the transition from yellow to pink occurs. These genomic clines are steeper than those expected under neutrality, suggesting that many of these *loci* are either under selection or physically linked to selected genes, most likely involved in pigment biosynthesis. The congruence between phenotypic and genotypic cline centers thus strongly supports the existence of a cryptic hybrid zone, in which gene flow between differentiated morphs is occurring, but is restricted and influenced by selection. Importantly the lack of population with both morphs reinforces the idea that this hybrid zone is not obvious in the landscape but is clearly detectable through integrated genetic and phenotypic analyses. This hybrid zone also provides a powerful framework to investigate the genetic basis of flower colour divergence. Cline-based approaches, combined with population genomics and transcriptomics, allowed us to identify candidate genes potentially underlying the observed phenotypic variation.

### On the genetic basis of flower colour variation in *P. comosa*

Hybrid zones are powerful natural systems to uncover the genetic basis of trait variation and its evolutionary implications. In the case of *P. comosa*, this approach is particularly valuable, as the species presents several challenges: it has a large genome (>2.5 Gbp), and its hemiparasitic lifestyle makes cultivation and experiments under controlled conditions difficult. Despite these constraints, our integrative approach enabled us to identify promising candidate genes involved in flower colour divergence.

Using nGBS, which sampled only a small fraction (that we estimated here at 0.22%) of the genome, we detected several outlier SNPs associated with strong allele frequency clines centered around the hybrid zone. Notably, among these candidates, we consistently identified Dihydroflavonol-4-reductase (DFR) through both AMOVA and cline analyses. DFR plays a key role in the anthocyanin biosynthesis pathway. Another promising candidate, also revealed by both methods, is a UDP-glycosyltransferase, which catalyzes glycosylation of flavonols such as quercetin and kaempferol. Parallel GWAS analyses uncovered an additional UDP-glycosyltransferase from a distinct genomic region, similar to a 7-deoxyloganetic acid glucosyltransferase-like protein, also likely involved in pigment biosynthesis. Interestingly, the geographic center of the strongest clines in both phenotypic traits and candidate SNPs coincides with the location of the well-studied hybrid zone between yellow and pink-flowered populations of *Antirrhinum majus* in the eastern Pyrenees (Whibley et al. 2006), suggesting that similar ecological or historical factors may shape flower colour transitions across unrelated plant taxa in this region.

To test the functional relevance of these candidates, we conducted RNA-seq analyses comparing pink and yellow individuals. DFR was confirmed as differentially expressed: strongly overexpressed in pink morphs and underexpressed in yellow ones. In addition, several other genes, not found with population genomic approaches, showed differential expression between morphs, including Flavonol synthase (FLS) which catalyzes the formation of flavonols from dihydroflavonols, Flavonoid 3’-hydroxylase (F3’H), Flavanone 3-hydroxylase (F3H), Anthocyanidin 3-O-glucosyltransferase (BZ1) and Anthocyanidin 3-O-glucoside 2’’’-O-xylosyltransferase (UGT79B1). While these genes were not detected based on the RAD-seq dataset, likely due to limited genome coverage, they are commonly associated with anthocyanin pigmentation in other species such as miniature roses (Lu et al. 2021) and this support the complementarity of the two approaches implemented in this study.

Although *P. comosa* displays a seemingly simple flower colour dimorphism, these results point to a complex genetic architecture, likely involving a few major *loci* and multiple secondary genes, as has also been shown in *Antirrhinum majus* (Bradley et al. 2017; Bradley et al. 2025; Field et al. 2025; Richardson et al. 2025). Among the most intriguing findings, we identified two tandemly duplicated copies of the FLS gene (FLS1 and FLS2) both composed of three similar exons. Notably, FLS1 harbours a large intron insertion and a non-synonymous mutation found only in pink morphs. This mutation substitutes a highly conserved arginine (R) with a glutamine (Q) near the catalytic site (Figure 6A, potentially altering enzyme function due to the difference in charge.

**Figure 6.**
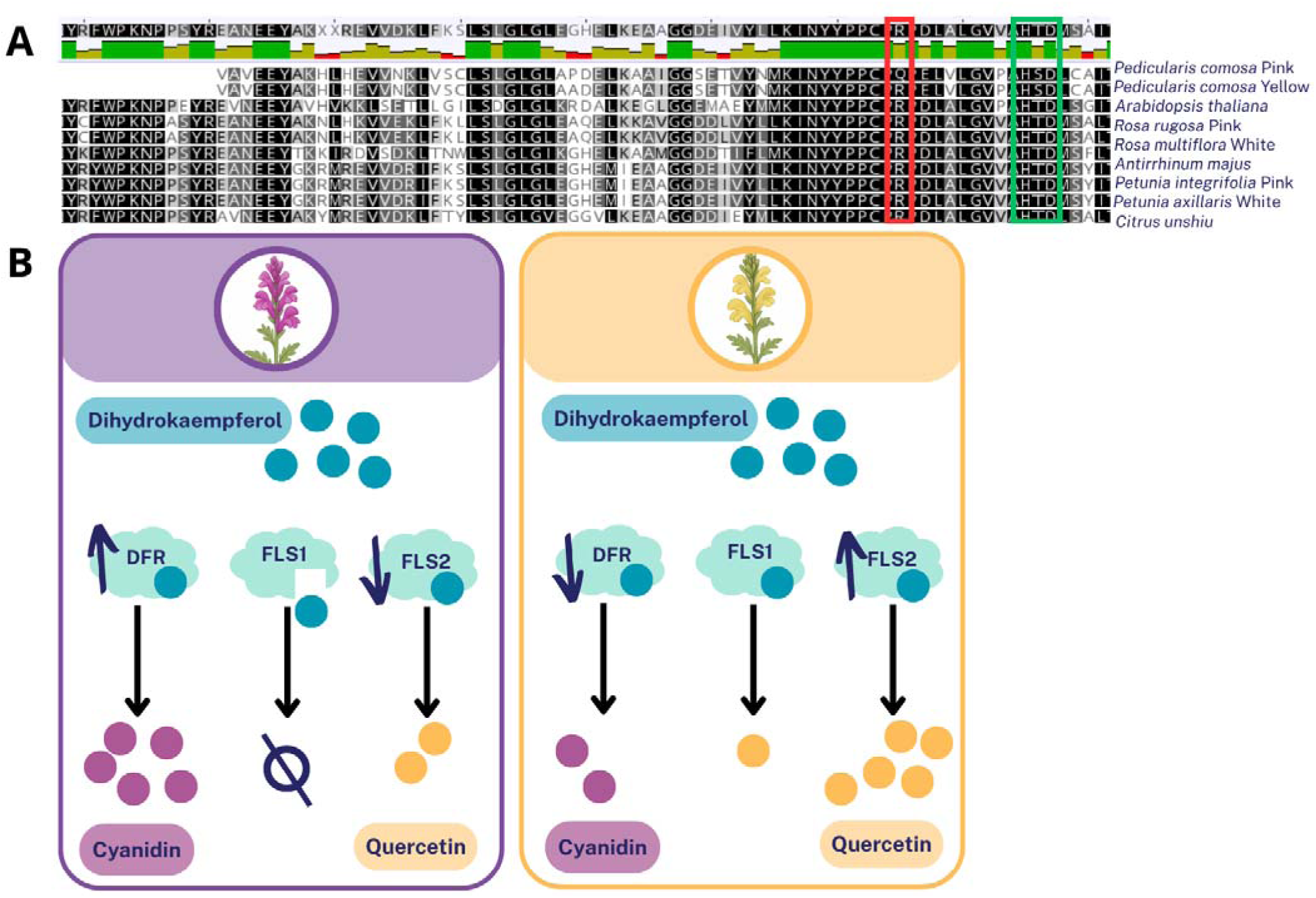
**A)** Protein sequence alignment of FLS1 from *P. comosa a*nd several phylogenetically distant plant species. Conserved regions are shown in black. The pink *P. comosa* morphotype displays a unique non-synonymous mutation (in red) substituting a conserved arginine (R) with a glutamine (Q) near the catalytic site (in green), potentially impacting enzyme function. **B)** Schematic summary of the model proposed to explain flower colour differentiation in *P. comosa*. The three key enzymes involved in pigment biosynthesis (DFR, FLS1 and FLS2) are shown. Arrows indicate their relative regulation in each floral morph – no arrow indicating no differential expression. Although FLS1 is not differentially expressed it carries a non-synonymous mutation that may affect its substrate affinity, suggesting competition among FLS1, FLS2 and DFR for shared metabolic intermediates, influencing the balance between anthocyanin (cyanidin) and flavonol (quercetin) production.

From these observations, we propose the following hypothesis (Figure 6B): in pink individuals, although FLS1 is not differentially expressed, the mutated form of the enzyme may still compete with DFR and FLS2 for their shared substrate, dihydroflavonol. This competition could reduce the synthesis of quercetin (a yellow pigment), while enabling higher levels of cyanidin (a pink pigment), due to the upregulation of DFR. In yellow individuals, the absence of the FLS1 mutation allows normal production of quercetin by both FLS1 and FLS2 while the low expression of DFR limits cyanidin accumulation, leading to yellow flowers.

These findings illustrate how even a simple phenotypic polymorphism can be explained by a complex, multilocus genetic basis. Although our study does not resolve the full molecular mechanisms, it establishes a strong foundation for further work, especially through whole-genome resequencing (WGS) and functional assays, to refine the genetic model of flower colour variation in *P. comosa*.

### On the eco-evolutionary causes of flower colour variation in *P. comosa*

Having identified several candidate genes likely involved in the divergence of floral pigmentation, including DFR, FLS and various glycosyltransferases, we now turn to the potential regulatory mechanisms and evolutionary drivers that may have shaped their expression and spatial distribution across the landscape.

At the molecular level, it has been recently supported that FLS and DFR enzymes compete for the same substrate, thereby influencing the balance between flavonol (yellow pigment) and anthocyanin (pink pigment) biosynthesis (Choudhary and Pucker 2024). However, their expression is likely regulated by upstream elements, including MYB transcription factors, such as ROSEA (a R2R3-MYB activator) and ELUTA (a repressor associated with a CACTA transposon), as described in Moss et al. (2024). In addition, small RNAs may also play a role in controlling pigment distribution. For example, Bradley et al. (2017) showed that small RNAs can restrict the accumulation of yellow aurones by targeting specific biosynthetic genes for silencing.

These regulatory pathways open the door to several evolutionary scenarios to explain the spatial distribution of flower colour morphs in *P. comosa*. One possibility (Supplementary Figure S9A) is that a recent genetic change promoting anthocyanin accumulation (e.g., through upregulation of DFR or altered competition with FLS) arose under selective pressures, such as the need to better tolerate warmer and drier conditions or to resist higher UV exposure, as reported in Grossenbacher et al. (2025), in the easternmost parts of the Pyrenees. This newly advantageous trait may now be spreading westward, including towards the Canigou massif. Such a dynamic could explain the partial mismatch between neutral genetic structure and the geographic distribution of flower colour morphs, with pigmentation changes outpacing genome-wide divergence.

An alternative scenario (Supplementary Figure S9B) is that pink-flowered populations in the Albères (1-East-Albères, 2-Pic Neulos, 3-Pic des Salines) and 4-Mariailles already existed before the Last Glacial Maximum (ca. 20,000 yrs ago) and maintained strong gene flow across the eastern range. After the LGM, gene exchange may have ceased, possibly due to the Tech River acting as a geographic barrier. Yellow morphs may then have expanded eastward from Southern/Western refugia, leading to a secondary contact zone between the Albères and the Canigou through the population of 4-Mariailles. Under this scenario, the hybrid zone would reflect ancient divergence and post-glacial range shifts, rather than recent adaptation.

At present, it remains difficult to discriminate between these scenarios, especially in the absence of clear information on pollinator assemblages, preferences, and behaviour in the Eastern Pyrenees. Indeed, little is known about pollinator identity or constancy in *P. comosa*, making it challenging to assess whether pollinator-mediated selection has played a role in shaping colour divergence.

Nonetheless, evidence from other *Pedicularis* species, particularly in the Hengduan Mountains of southwest China, where the genus reaches its highest diversity, provides informative comparisons. In this region, flower morphology often shows spatial structuring aligned with microhabitat rather than sharp species boundaries (Tang et al. 2023). Several species exhibit intraspecific variation in colour associated with parapatric distributions, such as *P. rex*, *P. kansuensis*, *P. batangensis* and *P. anas,* suggesting that colour transitions may evolve repeatedly from standing variation, possibly in response to local abiotic or biotic selection pressures.

In this context, the pink-yellow colour dimorphism in *P. comosa* may similarly reflect parallel ecological divergence. For example, anthocyanin-rich morphs may confer adaptive benefits under environmental stress (e.g. high UV, drought, high temperatures) or herbivory, as suggested in other plants (Li and Ahammed 2023). While speculative, this could explain the predominance of the pink morph in the easternmost, warmer and more UV-exposed parts of the range, and the westward expansion of this phenotype in the face of gene flow.

Finally, a less parsimonious but still plausible explanation is adaptive introgression where pink-flowered morphs of *P. comosa* may have acquired their pigmentation through historical gene flow from other pink-flowered *Pedicularis* species in sympatry. In the Pyrenees, several species such as *P. verticillata*, *P. rosea*, *P*. *palustris*, *P. kerneri*, *P. sylvatica*, *P. pyrenaica*, *P. mixta* share similar floral pigmentation. While we currently lack genomic evidence to support this hypothesis, it would be worth testing it with broader sampling and comparative phylogenomics.

### Conclusive remarks

Our results reveal a complex pattern of flower colour divergence in *P. comosa*, maintained by selection across a cryptic hybrid zone. The sharp clines and mismatch with neutral structure suggest an interplay of ecological and historical factors.

The Ibero-Pyrenean suture zone, formally recognised as such by Milá et al. (2013), stands out as a major hotspot of both inter- and intraspecific convergence. Acting as both a barrier and a refugium, the Pyrenees have influenced the genetic structure and hybridization patterns of numerous taxa (e.g., *Alytes*, *Cervus*, *Zootoca*, *Quercus*…), underscoring their evolutionary and conservation significance. Within this context, our findings position the eastern Pyrenees as a key area for preserving intraspecific diversity in *Pedicularis comosa*. We therefore advocate for the conservation of both flower colour morphs to safeguard the species’ full evolutionary potential.

## Materials and Methods

### Study sites and sample collection

We sampled 178 *Pedicularis comosa* individuals across 12 localities spanning the species’ distribution in the Eastern Pyrenees during the flowering seasons of 2021 and 2023 (Figure 1; Supplementary Note 2, Supplementary Table S4). For each plant, 1–2 cm² of leaf tissue was collected and preserved in 90% ethanol at 4 °C until DNA extraction. For transcriptomic and pigment analyses, we sampled pink and yellow flowers from a subset of 20 individuals at seven sites. Flowers were preserved in RNAlater (Fisher) for RNA extraction (stored at −20 °C), and additional floral tissues from the same individuals were dried on silica gel, then lyophilised (24 h) prior to LC–MS analyses. Calibrated photographs were taken for all individuals to quantify flower colour variation. Sampling in French nature reserves was conducted under permits issued by the *Direction Départementale des Territoires et de la Mer 66* (DDTM 66).

### Reference genome assembly and annotation

We used a combination of PacBio long-read sequencing and optical mapping to sequence and assemble a high-quality reference genome for *Pedicularis comosa* (Supplementary Note 1, Supplementary Table S1, S2, S3, Supplementary Figure S1). High-Molecular-Weight (HMW) DNA was extracted from a specimen of the species with pink flowers (21-Ped-135), which was collected from the centre of the study area (4-Mariailles). The assembly and hybrid scaffolding were done with HiFiasm v.0.15.5 (Cheng et al. 2021) and the *hybridScaffold* pipeline from Bionano, respectively. The structural annotation was performed with Eugene-EP v2.0.4 pipeline (Sallet et al. 2019; Carrere et al. 2023) with the help of floral transcriptomes of one pink-flowered and one yellow-flowered individual. All these procedures were carried out at the Centre National de Ressources Génomiques Végétales (CNRGV, Toulouse, France). In addition, Transposable Elements (TE) annotation was performed using both homology- and structure-based approaches, as previously described (Kim et al. 2024; Russo et al. 2024). Further details regarding the genome assembly and annotation can be found in Supplementary Note 1.

### Genotyping (nGBS) and SNP calling

We genotyped 156 *Pedicularis comosa* individuals using a normalised Genotyping by Sequencing (nGBS) approach (LGC Genomics GmbH, Berlin, Germany), a double digest RADseq-like method well suitable for large genomes. Individually barcoded libraries (2 × 250 bp) were sequenced on an Illumina NovaSeq 6000, yielding >1.5 million read pairs per sample. Reads were processed and mapped to the *P. comosa* subsp. *asparagoides* reference genome (assembled in this study) using Stacks v2.60 (Catchen et al. 2011; Catchen et al. 2013). After stringent filtering for coverage, minor allele frequency (≥0.05), and heterozygosity (≤0.7), the final dataset contained 28,729 SNPs for downstream population genomic, phylogenetic, and cline analyses (detailed pipeline in Supplementary Note 3).

### Phylogeny and population genomics analyses

We reconstructed a Maximum-Likelihood phylogeny of the 156 sampled *P. comosa* individuals using IQ-TREE 2 (Minh et al. 2020) based on 14,749 nGBS *loci* (28,729 SNPs). The best-fit model (GTR+F+I+G4) was selected by ModelFinder (Kalyaanamoorthy et al. 2017), and branch support was assessed with 1,000 ultrafast bootstrap replicates. Population structure and genetic diversity were analysed with PCA (adegenet; (Jombart and Ahmed 2011), *G*-statistics (Genodive v3.04; (Meirmans 2020), and Isolation by Distance (IBD) tested via a Mantel test (vegan; Oksanen 2017). We inferred ancestry coefficients with sNMF (LEA; (Frichot et al. 2014) and ADMIXTURE (Alexander et al. 2009) for *K* = 1–12, selecting the optimal *K*-values via cross-validation. FineRADstructure (Malinsky et al. 2018) was used to infer coancestry patterns. Complete parameter settings and filtering thresholds are provided in Supplementary Note 4.

### Phenotypic, genomic and geographic clines

To test whether the contact zone between pink- and yellow-flowered populations corresponds to a hybrid zone, we also fitted geographic clines for phenotypic and genomic data. Flower colour scores (cyan, magenta, yellow) were extracted from calibrated photographs for seven localities and used to fit phenotypic clines. Clines were fitted for hybrid indices (Q) from sNMF and ADMIXTURE (*K*=4), which best captured differentiation between the Albères and Canigou massifs. Sampling sites were projected onto a one-dimensional transect, and clines were fitted using HZAR (Derryberry et al. 2014) with a Metropolis–Hastings MCMC algorithm.

Genome-wide clines were also estimated for 28,729 SNPs using three equilibrium cline models (fixed center, free center, with/without tails). For each SNP, the best-fitting model was selected via AICc, and cline center and width were extracted. Steep clines were defined as those with allele frequency shifts (Δ > 0.6) across the transect, centers between 56–70 km, and widths ≤33 km. Twenty-eight candidate SNPs exhibited sharp allele frequency transitions (Figure□4B, Supplementary Table□S7, Supplementary Note 5).

### Analyses of introgression

To evaluate introgression among populations, we used Patterson’s D (ABBA–BABA) and the f4-ratio statistic (Patterson et al. 2012) as implemented in *Dsuite* v0.5 (Malinsky et al. 2021). These tests, unlike PCA or clustering approaches, formally assess admixture within an explicit phylogenetic framework. We ran *Dtrios* with the IQ-TREE topology (Figure 2B) as reference and 11-Capolat as the outgroup (the most divergent population), applying block jackknifing to assess significance.

We further used the f-branch approach to visualise admixture proportions across the tree, and *triangulaR* v0.0.1 (Wiens et al. 2025) to compute hybrid indices, interclass heterozygosity, and ancestry-informative markers, visualised via triangle plots. Parental populations were set as 1-East Albères and 7-Llo, representing the Albères and Canigou gene pools, respectively.

### Investigation of the genomic bases of colour variation

To identify *loci* contributing to genetic structuring and colour divergence, we combined two complementary genome scan approaches. First, we used BayPass v2.1 (Gautier 2015) to detect SNPs associated with flower colour, while explicitly accounting for population covariance due to shared demographic history. Populations were grouped into pink (1-East Albères, 2-Pic Neulos, 3-Pic des Salines, 4-Mariailles) and yellow morphs (5-Prats-de-Mollo, 6-Py-Mantet, 7-Llo, 8-Nohèdes, 9-Jujols, 10-Mijanès, 11-Capolat). Second, we performed an AMOVA-based genome scan (Genodive v3.04; Meirmans 2020) to identify *loci* showing the highest differentiation (*F*_CT_) between the two morph groups, partitioning variance hierarchically among morphs, populations, and individuals (Meirmans and Liu 2018). The 100 SNPs with the highest significant *F*_CT_ values were retained for further investigation. Third, we used genome-wide association studies (GWAS) to identify *loci* associated with flower colouration. Colour was considered both as qualitative trait (yellow *vs.* pink) and as quantitative traits based on three channel scores (Red, Magenta, and Yellow). Flower colour was quantified from standardised photographs by extracting values from the RGB (Red, Green, Blue) channels and the CMY (Cyan, Magenta, Yellow) colour space. For each population, mean channel values were calculated on one individual to provide objective measures of colour variation among populations. We applied three multi-locus models that control for population structure, kinship, linkage disequilibrium (LD), and co-variation among markers: the MLMM (multi-locus mixed linear model), the FarmCPU model (Fixed and random model Circulating Probability Unification), and the BLINK model (Bayesian-information and Linkage-disequilibrium Iteratively Nested Keyway). Simulation studies have shown these models to outperform simple mixed linear models (MLM) and general linear models (GLM) in terms of both statistical power and computational efficiency (Huang et al. 2019). All analyses were implemented in the R package GAPIT (Wang and Zhang 2021). *P*-values were adjusted using a false discovery rate (FDR) procedure (Benjamini and Hochberg 1995) as implemented in GAPIT, and quantile-quantile (QQ) plots were used to evaluate the fit between observed and expected *p*-value distributions.

### Transcriptome analysis

Total RNA was extracted from floral tissues using the Qiagen RNeasy Plant Mini Kit, quantified with Qubit 4.0, and checked for integrity on an Agilent 5600 Fragment Analyzer. Libraries were prepared with the NEBNext Ultra II RNA Library Prep Kit (New England Biolabs), following poly(A) selection, mRNA fragmentation (15□min, 94□°C), cDNA synthesis, adapter ligation, and PCR enrichment. Libraries were validated, pooled, and sequenced (2□×□150□bp, NovaSeq 6000, S4 lane). Raw reads were quality-checked (FastQC v0.11.9, (Andrews 2015), trimmed (Trimmomatic v0.39, (Bolger et al. 2014), and aligned to the *P. comosa* subsp. *asparagoides* genome using HISAT2 v2.2.1 (Kim et al. 2019). Read counts per gene were obtained with HTSeq-count v2.0.2 and normalised for differential expression analyses in DESeq2 (Love et al. 2014). Differential expression was assessed with Wald tests, and *p*-values were adjusted for multiple testing using the Benjamini-Hochberg procedure (FDR). Genes were considered significantly differentially expressed when *p*-adj < 0.05 and |log2FC| > 2 (details are provided in Supplementary Note 6).

### Pigment analysis

Lyophilised floral tissues were extracted with 80:20 MeOH/H₂O containing 1% acetic acid, agitated at 4□°C for 24□h, centrifuged, filtered (PTFE, 0.2□µm), and analysed by UHPLC-DAD-MS (Q-Exactive Plus Orbitrap, Thermo Fisher). Chromatographic separation was performed on a Luna Omega Polar C18 column (100□×□2.1□mm, 1.6□µm, Phenomenex) using a binary gradient (MP.A: acetonitrile□+□0.1% FA; MP.B: water□+□0.1% FA) at 0.4□mL□min⁻¹. UV detection spanned 200–800□nm; mass spectra were acquired in positive and negative electrospray modes (100–1500□*m/z*, MS resolution 70,000).

Raw MS data were converted to mzXML (ProteoWizard v3.0; (Chambers et al. 2012) and processed in MZmine v2.53 (Schmid et al. 2023) for peak detection, alignment, and gap filling. Molecular networks were generated using GNPS workflows (cosine score□>□0.7, ≥6 matched peaks; https://gnps.ucsd.edu/ProteoSAFe/static/gnps-splash.jsp) visualised in Cytoscape v3.8.2 (Shannon et al. 2003), and annotated using Sirius v5.6.2 (Dührkop et al. 2019) and plant metabolite databases. UV-Vis data were combined with MS/MS annotations to confirm compound identities.

## Supporting information

Supp_Mat

## Acknowledgements

This study was supported by the interreg-POCTEFA program (FLORALAB Project EFA294/19 and FLORALAB+ EFA 024/01) and a side project allocation grant from the “École Universitaire de Recherche (EUR)” TULIP-GS (ANR-18-EURE-0019) to PS. This work is set within the framework of the “Laboratoires d’Excellences (LABEX)” TULIP [ANR-10-LABX-41]. PS, AG and JAMB were also supported by an Agence Nationale pour la Recherche Jeune Chercheur Jeune Chercheuse (ANR JCJC) grant to JAMB, grant number ANR-21-CE02-0022-01. We would also like to express our gratitude to everyone who participated in the surveys : Joseph Garrigue, Diane Sorel, Christophe Hurson, Cécile Brousseau, Artur Lluent, Charles Esther, Bernard Latour, Dominique Aubert, Anne De Bures, Jean-André Magdalou, Léa Jugnet, Catherine André and Louis Boulesteix, Antoine Sénac, Kimberley Goudédranche as well as students of the Master 1 Biodiversité Écologie & Évolution (2020-2021): Paula Voegele, Magdalena Voisin-Baenitz, Noëline Amilhat, Ève Longépé, Hugo Bellavoir, Mélina Wyss, Johanna Stéfani, Raphaël Rousseau, Arthur Sinel, Camille Peltier, Mérédith Salémi, Amélie Hienne, Alexandra Marques, Camille Château, Claire Garcia, Joan Kolasa, Laurie Gaillard, Maxime Alberto and Marine Cuvilliez.

## Data availability statement

Sequencing data have been submitted to the Sequence Read Archive (SRA) under Study with primary accession n°PRJEB15465741.

