## Supplementary material for "A cryptic hybrid zone reveals the genomic basis of flower colour variation in a plant with a large and complex genome": Supp_Mat

^5^ Association Charles Flahault, Toulouges, France

^6^ C. Barcelona, Berga, Spain

This PDF file includes :

Supplementary Notes 1 - 15

Supplementary Tables 15 - 43

Supplementary Figures 43 - 51

**Supplementary Notes**

**Supplementary Note 1 : Reference genome sequencing, assembly and annotation**

***High Molecular Weight (HMW) DNA extraction***

High Molecular Weight (HMW) DNA of *Pedicularis comosa* was extracted from individual 21-Ped-135 (4-Mariailles) frozen stems with buds using QIAGEN Genomic-tips 500/G kit (Qiagen, MD, USA). We followed the tissue protocol extraction. Briefly, 1.5 g of young plant material were grounded in liquid nitrogen with mortar and pestle. After 2h of lysis and one centrifugation step, DNA was first immobilized on the column. After several washing steps, DNA was eluted from the column, then desalted and concentrated by Isopropyl alcohol precipitation. A final wash in 70% ethanol was performed before resuspending the DNA in EB buffer. Analyses of DNA quantity and quality were performed using NanoDrop and Qubit (Thermo Fisher Scientific, MA, USA). DNA integrity was also assessed using the Agilent FP-1002 Genomic DNA 165 kb on the Femto Pulse system (Agilent, CA, USA).

***PacBio Hifi library preparation and sequencing***

Hifi libraries were constructed using SMRTbell® Template Prep kit 2.0 (Pacific Biosciences, Menlo Park, CA, USA) according to PacBio recommendations (SMRTbell® express template prep kit 2.0 - PN: 100-938-900). HMW DNA samples were first purified with 1X Agencourt AMPure XP beads (Beckman Coulter, Inc, CA USA), and sheared with Megaruptor 3 (Diagenode, Liège, BELGIUM) at an average size of 20 kb. After End repair, A-tailing and ligation of SMRTbell adapter, the library was size selected on BluePippin System (Sage Science, MA, USA) at range size of 10-50 kb. The size and concentration of libraries were assessed using the Agilent FP-1002 Genomic DNA 165 kb on the Femto Pulse system and the Qubit dsDNA HS reagents Assay kit. Sequencing primer v5 and Sequel® II DNA Polymerase 2.2 were annealed and bound, respectively to the SMRTbell libraries. Each library was loaded on 2 SMRTcell 8M at an on-plate concentration of 90pM. Sequencing was performed on the Sequel® II system at Gentyane Genomic Platform (INRAE Clermont-Ferrand, France) with Sequel® II Sequencing kit 3.0, a run movie time of 30 hours with an Adaptive Loading target (P1 + P2) at 0.75.

***Genome assembly***

109 Gb of High Fidelity (HiFi) reads were produced with PacBio Sequel II system on 4 SMRTCells and were assembled using HiFiasm (v0.15.5, Cheng et al. 2021). Hifiasm is able to produce a primary assembly and alternative assembly (incomplete alternative assembly consisting of haplotigs under heterozygous regions). We obtained a high-quality primary assembly of 2.87 Gb in 401 contigs with 22 Mb of N50 and an alternative assembly of 2.25 Gb.To assess the completeness and quality of the genome, we used the Benchmarking Universal Single-Copy Orthologs (BUSCO) pipeline with the viridiplantae database (Simão et al. 2015). We obtained a 98.4% complete BUSCO score on the primary assembly. In addition, we perform k-mer analysis to quality control the dataset using Jellyfish tool (Marçais and Kingsford 2011) and the assemblies using module “comp” of the k-mer Analysis Toolkit (Mapleson et al. 2017).

***uHMW DNA extraction and scaffolding with optical maps***

uHMW DNA was purified from 1g of apex flowers according to Bionano Prep Plant Tissue DNA Isolation Base Protocol (30068 - Bionano Genomics) with the following specifications and modifications. Plant material was ground in liquid nitrogen and then disrupted using a Tissue Ruptor grinder (QIAGEN, Redwood City, CA, USA) in homogenization buffer containing spermine, spermidine and beta-mercaptoethanol. Nuclei were washed, purified using a density gradient and then embedded in agarose plugs. After overnight proteinase K (QIAGEN, Redwood City, CA, USA) digestion in Lysis Buffer (Bionano Genomics) and 1-hour treatment with RNAse A (Qiagen), plugs were washed and solubilized with AGARase enzyme (ThermoFisher Scientific). A dialysis step was performed in TE Buffer (ThermoFisher Scientific) for 45 minutes to purify DNA from any residues. The DNA samples were quantified by using the Qubit dsDNA BR Assay (Invitrogen). The presence of mega base size DNA was visualized by pulsed field gel electrophoresis (PFGE).

Labelling and staining of the uHMW DNA were performed according to the Direct Label and Stain (DLS) protocol (30206 - Bionano Genomics). Briefly, labelling was performed by incubating 750 ng genomic DNA with 1× DLE-1 Enzyme for 2 hours in the presence of 1× DL-Green and 1× DLE-1 Buffer. Following proteinase K digestion and DL-Green clean-up by membrane adsorption, the DNA backbone was stained by mixing the labelled DNA with DNA Stain solution in the presence of 1×Flow Buffer and 1× DTT and incubating 1h at room temperature. The DLS DNA concentration was measured with the Qubit dsDNA HS Assay (Invitrogen, Carlsbad, CA, USA).

Labelled and stained DNA was loaded on 1 Saphyr chip and was run on the BNG Saphyr System according to the Saphyr System User Guide (30247 – Bionano Genomics). Digitalized labelled DNA molecules were assembled to optical maps using the BNG Access software (solve version 3.5).

***Hybrid scaffolding***

The optical map assembly reach 4.8 Gb and 22.7 Mb of N50. This large assembly size indicates that both allelic maps of this heterozygous diploid genome were obtained. To realize the scaffolding, we had to purge the duplicated maps corresponding to both alleles. This method was performed using an in-house script (described in the supplemental material of this article Rimbault et al. 2023) and allow to reduce the optical map from 4.7 Gb to 2.65 Gb (87 maps and 127 Mb of N50). This reduced optical map was used to scaffold the primary assembly with the hybridScaffold pipeline (<https://bionano.com/wp-content/uploads/2023/01/30073-Bionano-Solve-Theory-of-Operation-Hybrid-Scaffold.pdf>; solve version 3.6). We obtained a hybrid scaffold genome of 2.8 Gb with only 48 scaffolds and 161 Mb of N50.

***Structural annotation***

Structural gene annotation was performed using the Eugene-EP v2.0.4 pipeline (Sallet et al. 2019; Carrere et al. 2023) which combined three training sets for structural annotation, including the RNA-Seq data from *Pedicularis* genotypes and four different protein databases: SwissProt, TrEMBL Plants, and predicted proteins from the *Brachypodium distachyon* and *Arabidopsis thaliana* reference genome.

***Transposable Elements annotation***

The annotation of transposable elements (TEs) was carried out using a combination of homology-based and structure-based strategies. In the first step, we aligned TE protein datasets from REXdb (Neumann et al. 2019) against the *P. comosa* assembled genome using fastx32 (Pearson 1999) with an e-value cutoff of 1e-6. Genomic intervals corresponding to overlapping hits of same TE superfamily were subsequently merged with Bedtools merge (Quinlan and Hall 2010) accounting for strand orientation (-s option). For each TE superfamily, the corresponding nucleotide sequences were extracted in FASTA format.

To cluster related elements, we performed an all-against-all BLASTn with a e-value threshold of 1e-50, followed by clustering using SiLiX (Miele et al. 2011) with a minimum requirement of 80% sequence identity across at least 80% of alignment coverage. At this stage, only coding regions of TEs were detected, and boundaries remained undefined. To refine these, we extracted 10 kb flanking regions for each paralog and aligned them with pblat, removing regions without homologous support across paralogs. Multiple sequence alignments were then conducted with MAFFT, and family-level consensus sequences were constructed.

In parallel, LTRharvest (Ellinghaus et al. 2008) was applied with adjusted parameters (-xdrop 37, -motif tgca, -motifmis 1, -minlenltr 100, -maxlenltr 3000, -mintsd 2) to identify additional LTR retrotransposons. Paralogous sequences recovered by this approach were clustered with SiLiX. Furthermore, MITEs were detected using MITE-tracker under default parameters (Han and Wessler 2010).

All TE families obtained from the in-house pipeline, LTRharvest, and MITE-tracker were merged, and redundancies were removed, yielding a final set of 2269 non-redundant consensus TE sequences. This library comprised 1047 Class I and 1222 Class II elements.

**Supplementary Note 2: Study sites and sample collection**

To investigate Differential Gene Expression between pink- and yellow-flowered individuals of *P. comosa*, we re-sampled (in 2023) the following populations: 2-Pic Neolous, 3-Pic des Salines, 4-Mariailles, 6-Py-Mantet, 7-Llo and 10-Mijanès we also collected individuals from a population in the Massif Central (Hures-la-Parade, Lozère, Occitanie, France) to be considered as outgroup. We visited but could not find any flowering individual of *P. comosa* in the following localities: 1-East Albères, 5-Prats-de-Mollo, 8-Nohèdes, 9-Jujols probably due to serious drought. Only the locality 11-Capolat could not be visited. To quantify color variation, we used a Canon EOS 70D camera and EF-S 15-85 mm F/3.5-5.6 IS USM lens to take calibrated pictures. On each picture, a grey chart allowed us to adjust the white balance during the post-processing.

**Supplementary Note 3: Genotyping (nGBS and SNP calling)**

To sample the large genome of Pedicularis comosa (>2.5 Gbp) across a substantial number of individuals, we used a **normalized GenotypingbySequencing (nGBS)** protocol (LGC Genomics GmbH, Berlin), a doubledigest RADseqlike method. Genomic DNA was digested with **PstI** and **ApeKI**, followed by adapter ligation and a normalization step to limit overrepresentation of repetitive elements. Individually barcoded libraries (n = 158) were sequenced in paired-end mode (2 × 250 bp) on an Illumina NovaSeq 6000, producing ≥1.5 million read pairs per sample.

Raw reads were processed with **Stacks v2.60** (Catchen et al. 2013) as follows:

- **process_radtags** was used to demultiplex, filter, and clean reads.
- Reads were aligned to the P. comosa subsp. asparagoides draft genome using **bwa v0.7.17** (Li 2013).
- **ref_map.pl** and **populations** modules were run to assemble loci and export genotypes.

***Filtering criteria:***

Loci genotyped in ≥80% of individuals per population (-r 80) and present in ≥5 populations (-p 5). Sites with minor allele frequency <0.05 (--min-maf 0.05) were removed. Loci with observed heterozygosity >0.7 (--max-obs-het 0.7) were discarded to minimize inclusion of paralogs. Two individuals (21‑Ped‑57 and 21‑Ped‑62) with excessive missing data were excluded. Only a single representative SNP per overlapping site was retained for admixture analysis (--ordered-export). The final dataset comprised **28,271 high-quality SNPs**, exported in multiple formats (VCF, STRUCTURE, phylip, genepop, finerad, and hzar) for population structure, phylogenetic, and cline analyses.

While SNP calling was primarily based on the P. comosa reference genome, we also performed **de novo** variant calling with **Stacks v2.60** (Catchen et al. 2011; Catchen et al. 2013) to validate results. Pipeline parameters were optimized on a subset of 12 representative individuals (covering geographic and sequencing diversity). Following recommendations (Paris et al. 2017; Rochette and Catchen 2017), we fixed m = 3 (minimum reads per allele) and varied M and n (mismatches allowed within and between loci), ultimately selecting -m 3, -M 6, -n 6 as the optimal combination to maximize assembled and polymorphic loci. This configuration was then applied to the full dataset.

**Supplementary Note 4: Phylogeny and population genomics analyses**

Phylogenetic relationships among the 156 sampled P. comosa individuals were inferred with IQ-TREE 2 (Minh et al. 2020) using 14,749 concatenated nGBS loci (28,271 SNPs). The substitution model GTR+F+I+G4 was selected by ModelFinder (Kalyaanamoorthy et al. 2017) based on the Bayesian Information Criterion (BIC). Node support was assessed with 1,000 ultrafast bootstrap replicates (-B 1000). The resulting unrooted tree was visualized with FigTree v1.4.4 (Rambaut 2009). Population genetic diversity was quantified as expected (*H*_E_) and observed heterozygosity (*H*_O_), total allelic richness (*A*), and private alleles (*A_p_*) using Genodive v3.04 (Meirmans 2020). Deviations from panmixia were measured with *G*_IS_, and significance tested via 10,000 permutations. Pairwise and overall *G*_ST_ (Nei 1987) were computed similarly. We performed a PCA (adegenet; (Jombart and Ahmed 2011) on the 156 × 28,271 SNP matrix (genind format) to visualize genetic structure. Isolation by Distance (IBD) was assessed by correlating ln(*F*_ST_/(1–*F*_ST_)) with ln-geographic distances (Rousset 1997), significance tested by Mantel test (Pearson, 999 permutations) using vegan v2.5-7 (Oksanen 2017). Population structure was inferred with sNMF (LEA v3; (Frichot et al. 2014; Frichot and François 2015) and ADMIXTURE v1.3 (Alexander et al. 2009) for *K* = 1–12, each run with 10 replicates. Optimal *K* was selected via cross-entropy (sNMF) and cross-validation error (ADMIXTURE). FineRADstructure (Malinsky et al. 2018) was used to infer coancestry matrices based on haplotype sharing. Membership proportions to each cluster were averaged by population along the transect for cline analyses.

**Supplementary Note 5: Phenotypic, genomic, and SNP-based clines**

We tested whether the contact zone between pink- and yellow-flowered P. comosa populations corresponds to a hybrid zone by fitting clines for phenotypic traits, genomic hybrid indices, and individual SNPs.

***Phenotypic clines***

Colour scores (cyan, magenta, yellow; CMY system) were extracted from calibrated photographs of flowers from seven localities (1-East Albères, 2-Pic Neulos, 3-Pic des Salines, 4-Mariailles, 6-Py-Mantet, 7-Llo, and 10-Mijanès). Due to flower wilting and drought-related absence of blooming, the remaining localities could not be scored.

***Genomic clines (hybrid indices)***

Hybrid indices (Q) were computed from ancestry proportions inferred by sNMF (K=4) and ADMIXTURE (K=4), chosen as these K values best resolved differentiation between the Albères and Canigou massifs. Sampling sites were collapsed to a 1D transect by linear regression of geographic coordinates, and distances (km) were calculated along the transect. Hybrid indices and colour scores were fitted to equilibrium cline models using HZAR v0.2-5 (Derryberry et al. 2014) with Metropolis–Hastings MCMC (scaling fixed; no tails).

***Genome-wide SNP clines***

We fitted clines to 28,271 SNPs (Stacks “hzar” output) using three equilibrium models: (i) fixed center, no tails; (ii) free center, no tails; and (iii) free center, both tails. The best-fitting model for each SNP was chosen by AICc. Maximum-likelihood estimates of cline center and width were extracted (Supplementary Figures S7–S8). SNPs were retained as candidates if they met all criteria: Δ allele frequency >0.6 between the westernmost (11-Capolat) and easternmost (1-East Albères) populations, cline centers 56–70 km, and widths ≤33 km. Twenty-eight SNPs with steep geographic clines were retained (Figure 4B, Supplementary Table S7).

**Supplementary Note 6: Transcriptome sequencing and differential expression analysis**

***RNA extraction and library preparation***

Total RNA was extracted from *P. comosa* floral tissues using the Qiagen RNeasy Plant Mini Kit (Qiagen, Hilden, Germany) following manufacturer’s instructions. RNA concentration was quantified with a Qubit 4.0 Fluorometer (Invitrogen, Carlsbad, USA) and integrity assessed on an Agilent 5600 Fragment Analyzer (Agilent Technologies). RNA-seq libraries were constructed using the NEBNext Ultra II RNA Library Prep Kit for Illumina (New England Biolabs, Ipswich, USA). Poly(A)+ mRNAs were enriched using Oligod(T) beads, fragmented (15 min at 94 °C), and converted to cDNA (first and second strand synthesis). Following end repair, 3′ adenylation, and adapter ligation, libraries were indexed and enriched via limited-cycle PCR. Final libraries were validated on the Fragment Analyzer and quantified via Qubit.

***Sequencing***
Validated libraries were multiplexed, clustered, and sequenced on an Illumina NovaSeq 6000 (S4 lane, paired-end 2 × 150 bp). Base calling was performed using NovaSeq Control Software (NCS), and raw bcl files converted to FASTQ using bcl2fastq v2.20 (allowing one mismatch in index identification).

***Read processing and mapping***

Reads were inspected with FastQC v0.11.9 (Andrews 2015) and trimmed using Trimmomatic v0.39 (Bolger et al. 2014) to remove adapters, low-quality bases (phred < 15), and reads <36 bp. Clean reads were aligned to the P. comosa subsp. asparagoides reference genome using HISAT2 v2.2.1 (Kim et al. 2019) with default parameters.

***Quantification and differential expression***

Read counts per gene were obtained with HTSeq-count v2.0.2 and normalized for differential expression analyses in DESeq2 (Love et al. 2014). The statistical design accounted for both batch effects and flower color morph (~ batch + condition), with the yellow morph set as the reference condition. Genes with fewer than 10 reads in at least eight samples (corresponding to the smallest group size) were filtered out prior to analysis. Normalization factors were estimated using DESeq2’s median-of-ratios method, and normalized counts were exported for downstream visualization. Differential expression was assessed with Wald tests, and p-values were adjusted for multiple testing using the Benjamini-Hochberg procedure (FDR). Genes were considered significantly differentially expressed when padj < 0.05 and |log2FC| > 2. To identify functional candidates, we compiled lists of genes involved in metabolic pathways of interest based on KEGG annotations: (i) pigment biosynthesis (phenylpropanoid, flavonoid, anthocyanin, isoflavonoid, flavone/flavonol, and carotenoid pathways; KEGG maps: 00940, 00941, 00942, 00943, 00944, 00906), and (ii) volatile/odor-related pathways (monoterpenes, sesquiterpenes, fatty acid derivatives, and volatile phenylpropanoids; KEGG maps: 0061, 0062, 00900, 00902, 00909, 00904). Normalized expression levels of candidate genes from these pathways were further explored through histogram-based visualizations to compare expression profiles between color morphs.

**Supplementary Note 7: Pigments analysis (LC-MS)**

***Metabolite extraction and UHPLC-HRMS profiling***

Lyophilized flowers (24 h) were extracted in 80:20 MeOH/H₂O with 1% acetic acid, agitated for 24 h at 4 °C, centrifuged (4500 rpm, 5 min, 10 °C), and filtered (Acrodisc H-PTFE, 0.2 µm). Extracts were diluted to 0.5 mg mL⁻¹ in MeOH and analyzed by UHPLC-DAD-MS (Q-Exactive Plus Orbitrap, Thermo Fisher Scientific). Separation used a Luna Omega Polar C18 column (100 × 2.1 mm, 1.6 µm; Phenomenex) with a binary gradient (MP.A: acetonitrile + 0.1% FA; MP.B: water + 0.1% FA): 0–1 min, 5% A; 1–26 min, linear ramp to 100% A; 26–30 min, 100% A; 30–32 min, ramp to 5% A; 32–35 min, 5% A, at 0.4 mL min⁻¹. UV spectra were recorded between 200–800 nm, and MS acquisition was performed in both positive and negative electrospray ionization modes over 100–1500 m/z (MS resolution 70,000; MS/MS 17,500 at 200 Da). The five most intense precursor ions were fragmented (20/30/40 eV), with isotopic exclusion enabled to improve network quality.

***Data processing***

Raw UHPLC-HRMS files were converted to mzXML (ProteoWizard v3.0.19202; (Chambers et al. 2012) and processed with MZmine v2.53 (Schmid et al. 2023). Noise thresholds were set to 1000 (mass detection) and 2000 (chromatogram building), with m/z tolerance of 0.005 Da (10 ppm) and retention time (RT) tolerance of 0.2 min. Peaks were deconvoluted using the local minimum search algorithm (S/N ≥ 10, RT range 0.05–0.5 min) and deisotoped using isotopic peak grouping. Alignment was performed with the Join Aligner (10 ppm m/z tolerance, 0.5 min RT), followed by gap filling.

***Molecular networking and compound annotation***

Molecular networks were generated in both ionization modes via GNPS ([https://gnps.ucsd.edu](https://gnps.ucsd.edu/)). MS/MS spectra were window-filtered (top 6 peaks per ±50 Da window) and clustered with MSCluster (parent m/z tolerance 0.02 Da; fragment tolerance 0.02 Da). Edges were retained if nodes were within each other’s top 10 matches, with cosine scores > 0.7 and ≥6 matched peaks. Library matches used GNPS and CASMI spectral libraries under identical filtering criteria. Networks were visualized in Cytoscape v3.8.2 (AllegroLayout; (Shannon et al. 2003), merged across ionization modes, and curated to remove background or processing artifacts. Formula predictions were performed in Sirius v5.6.2 (C/H/O only, 5 ppm mass tolerance, 20% isotopic tolerance; (Dührkop et al. 2019). Chemical classes were retained for matches with scores > 5, and structures were accepted for similarity ≥ 70%. UV-Vis spectra were integrated with MS/MS data to confirm the identity of major anthocyanin and flavonoid derivatives.

***Identification and characterization of anthocyanin pigments***

LC–MS profiling revealed two dominant anthocyanin-related metabolites associated with floral pigmentation differences in *Pedicularis comosa*. In positive ionization mode, two major ions were detected at m/z 787.19 (retention time tₙ = 5.29 min) and m/z 595.17 (tₙ = 5.64 min). The ion at m/z 787.19 was detected in both floral morphs but was significantly more abundant in pink flowers, whereas the ion at m/z 595.17 was detected exclusively in pink morphs and was absent from all yellow-flowered individuals, including across all detected isomers (Wilcoxon tests). Both compounds exhibited strong UV–visible absorbance maxima around 520 nm, consistent with anthocyanidin chromophores. Molecular networking analyses revealed that the m/z 787.19 feature clustered with compounds sharing a diagnostic fragment at m/z 303.04, corresponding to a delphinidin aglycone. By contrast, the m/z 595.17 feature clustered with ions producing a fragment at m/z 287.05, characteristic of cyanidin-based pigments. MS/MS fragmentation of the m/z 595.17 ion yielded a prominent fragment at m/z 449.11, consistent with sequential neutral losses of 162.05 Da and 146.05 Da. The UV spectrum of this compound showed a characteristic shoulder around 332 nm, indicating that the 162 Da loss likely corresponds to a coumaroyl moiety rather than a hexose, while the 146 Da loss is consistent with rhamnose ([M+H–H₂O]). Based on these combined MS/MS and UV–visible features, this compound was annotated as a cyanidin–coumaroyl–rhamnoside, a cyanidin-derived acylated anthocyanin. For the m/z 787.19 ion, MS/MS spectra revealed a major fragment at m/z 479.08, corresponding to the sequential loss of 176.03 Da and 308.11 Da. The 176 Da loss is consistent with a hexose moiety, whereas the 308 Da loss corresponds to a rutinose disaccharide. Together with the delphinidin-derived aglycone signature, these data support the annotation of this metabolite as a delphinidin-derived rutinoside.

**Supplementary Tables**

**Supplementary Table S1 :** Features of genome assembly of *P. comosa asparagoides*

| Pacbio data | 30.49 G |
| --- | --- |
| Estimated genome size | 2.72 Gb |
| Total length of scaffolds | 2814.098 Mb |
| Number of scaffolds | 48 |
| Scaffold N50 | 161.094 Mb |
| Contig N50 | 22,121,117 bp |
| Assembly completedness estimation (BUSCO) | 98.4%, n=425 |

**Supplementary Table S2 :** Gene structure Prediction statistics of *P. comosa asparagoides*

| Protein coding genes | Number of protein coding genes | 48030 |
| --- | --- | --- |
|  | Mean gene length (bp) | 2394.03 |
|  | Coding nucleotides (bp) | 46930949 |
|  | Percent genes with introns | 71 |
|  | Percent genes with five UTR | 45 |
|  | Percent genes with three UTR | 45 |
|  | **EXONS** |  |
|  | Mean number per gene | 4.02 |
|  | Mean length (bp) | 311.53 |
|  | GC per cent | 43.7 |
|  | **INTRONS** |  |
|  | Mean number per gene | 3.02 |
|  | Mean length (bp) | 377.08 |
|  | GC per cent | 31.46 |
|  | **CDS** |  |
|  | Mean length (bp) | 977.12 |
|  | Min length (bp) | 123 |
|  | Max length (bp) | 15279 |
|  | GC per cent | 45.75 |
|  | **five prime UTR** |  |
|  | Mean length (bp) | 270.82 |
|  | GC per cent | 38.14 |
|  | **three prime UTR** |  |
|  | Mean length (bp) | 342.66 |
|  | GC per cent | 35.15 |
| Non protein coding genes | Number of non protein coding genes | 45878 |
|  | Mean ncRNA gene length (bp) | 380.47 |
|  | Min length (bp) | 45 |
|  | Max length (bp) | 31538 |
|  | GC per cent | 51.34 |
|  | Percent ncRNAgenes with introns | 0 |
|  | Mean exon number per ncRNA gene | 1 |
| Intergenic | Mean length (bp) | 28648.35 |
|  | GC per cent | 37.87 |

**Supplementary Table S3:** TE annotation statistics of *P. comosa asparagoides*

| Wicker code | Class | Order | Superfamily | Total Length (bp) | Genome % | % of Total TEs |
| --- | --- | --- | --- | --- | --- | --- |
| RLC | Class I | LTR | Copia | 344019405 | 12,22 | 17,01 |
| RLG | Class I | LTR | Gypsy | 987010636 | 35,07 | 48,81 |
| RLX | Class I | LTR | NA | 210198481 | 7,47 | 10,39 |
| RIL | Class I | LINEs | NA | 108710505 | 3,86 | 5,38 |
| DTC | Class II | TIR | CACTA | 61611737 | 2,19 | 3,05 |
| DTM | Class II | TIR | Mutator | 62837363 | 2,23 | 3,11 |
| DTA | Class II | TIR | hAT | 42452054 | 1,51 | 2,1 |
| DTH | Class II | TIR | PIF-Harbinger | 48893781 | 1,74 | 2,42 |
| DTT | Class II | TIR | Mariner | 4524739 | 0,16 | 0,22 |
| DHH | Class II | RC | Helitron | 81312095 | 2,89 | 4,02 |
| DXX | Class II | TIR | MITEs | 70623833 | 2,51 | 3,49 |
| Total |  |  |  | 2022194630 | 71,8594151 | 100 |

**Supplementary Table S4** : Individual ID, locality, geographic coordinates: longitude and latitude (in °, WGS84), altitude, taxonomy, sample type: DNA, RNA, calibrated pictures and collector name(s)

Collector names: ADB : Anne DE BURRES AG : Anaïs GIBERT, BL : Bernard LATOUR, CB : Cécile BROUSSEAU, CH : Christophe HURSON, CQ : Céline QUELENNEC, DS : Diane SOREL, JAM : Jean-André MAGDALOU, JB: Joris BERTRAND, JG: Joseph GARRIGUES, JML : Jean-Marc LEWIN, LJ : Léa JUGNET, MM: Maria MARTIN, PA: Pere AYMERICH, PG : Pascal GAULTIER, PS : Pascaline SALVADO, VH : Valérie HINOUX.

| Individual_ID | Locality | Longitude | Latitude | Altitude | Species | RNA Sampling | Calibrated picture | Remark |
| --- | --- | --- | --- | --- | --- | --- | --- | --- |
| 21-Ped-01 | East_Alberes | 3.0094533 | 42.4676933 | 988 | *P. c. asparagoides* |  | x | Collectors VH, JB |
| 21-Ped-02 | East_Alberes | 3.0091301 | 42.4676567 | 992 | *P. c. asparagoides* |  |  | Collectors VH, JB |
| 21-Ped-03 | East_Alberes | 3.0306533 | 42.4744367 | 928 | *P. c. asparagoides* |  | x | Collectors VH, JB |
| 21-Ped-04 | East_Alberes | 3.0327667 | 42.4733983 | 899 | *P. c. asparagoides* |  | x | Collectors VH, JB |
| 21-Ped-05 | East_Alberes | 3.0341299 | 42.4735616 | 914 | *P. c. asparagoides* |  | x | Collectors VH, JB |
| 21-Ped-06 | East_Alberes | 3.0342334 | 42.4733650 | 905 | *P. c. asparagoides* |  | x | Collectors VH, JB |
| 21-Ped-07 | East_Alberes | 3.0379066 | 42.4736317 | 997 | *P. c. asparagoides* |  | x | Collectors VH, JB |
| 21-Ped-08 | East_Alberes | 3.0376901 | 42.4734067 | 982 | *P. c. asparagoides* |  |  | Collectors VH, JB |
| 21-Ped-09 | East_Alberes | 3.0264033 | 42.4718316 | 977 | *P. c. asparagoides* |  | x | Collectors VH, JB |
| 21-Ped-10 | East_Alberes | 3.0045150 | 42.4696766 | 1008 | *P. c. asparagoides* |  | x | Collectors VH, JB |
| 21-Ped-11 | East_Alberes | 3.0028500 | 42.4710201 | 1045 | *P. c. asparagoides* |  | x | Collectors VH, JB |
| 21-Ped-12 | East_Alberes | 3.0015934 | 42.4720383 | 1085 | *P. c. asparagoides* |  |  | Collectors VH, JB |
| 21-Ped-13 | East_Alberes | 3.0004783 | 42.4726884 | 1112 | *P. c. asparagoides* |  | x | Collectors VH, JB |
| 21-Ped-14 | East_Alberes | 2.9988884 | 42.4722834 | 1135 | *P. c. asparagoides* |  |  | Collectors VH, JB |
| 21-Ped-15 | East_Alberes | 2.998488 | 42.472447 | 1197 | *P. c. asparagoides* |  |  | Collectors VH, JB |
| 21-Ped-16 | Pic_Neolous | 2.9469367 | 42.4821717 | 1261 | *P. c. asparagoides* |  |  | Collectors VH, JB |
| 21-Ped-17 | Pic_Neolous | 2.9469617 | 42.4822367 | 1257 | *P. c. asparagoides* |  |  | Collectors VH, JB |
| 21-Ped-18 | Pic_Neolous | 2.9470601 | 42.4822267 | 1263 | *P. c. asparagoides* |  |  | Collectors VH, JB |
| 21-Ped-19 | Pic_Neolous | 2.9491601 | 42.4824267 | 1242 | *P. c. asparagoides* |  |  | Collectors VH, JB |
| 21-Ped-20 | Pic_Neolous | 2.9490367 | 42.4824401 | 1241 | *P. c. asparagoides* |  |  | Collectors VH, JB |
| 21-Ped-21 | Pic_Neolous | 2.9466700 | 42.4821234 | 1258 | *P. c. asparagoides* |  |  | Collectors VH, JB |
| 21-Ped-22 | Pic_Neolous | 2.9454667 | 42.4826483 | 1223 | *P. c. asparagoides* |  |  | Collectors VH, JB |
| 21-Ped-23 | Pic_Neolous | 2.9453067 | 42.4828384 | 1219 | *P. c. asparagoides* |  |  | Collectors VH, JB |
| 21-Ped-24 | Pic_Neolous | 2.9449867 | 42.4829234 | 1218 | *P. c. asparagoides* |  |  | Collectors VH, JB |
| 21-Ped-25 | Pic_Neolous | 2.9449716 | 42.4830300 | 1211 | *P. c. asparagoides* |  |  | Collectors VH, JB |
| 21-Ped-26 | Pic_Neolous | 2.9448866 | 42.4832550 | 1200 | *P. c. asparagoides* |  |  | Collectors VH, JB |
| 21-Ped-27 | Pic_Neolous | 2.9480550 | 42.4858867 | 1158 | *P. c. asparagoides* |  |  | Collectors VH, JB |
| 21-Ped-28 | Pic_Neolous | 2.9480951 | 42.4860367 | 1155 | *P. c. asparagoides* |  |  | Collectors VH, JB |
| 21-Ped-29 | Pic_Neolous | 2.9482516 | 42.4857883 | 1174 | *P. c. asparagoides* |  |  | Collectors VH, JB |
| 21-Ped-30 | Pic_Neolous | 2.9484550 | 42.4857900 | 1165 | *P. c. asparagoides* |  |  | Collectors VH, JB |
| 21-Ped-31 | Pic_Neolous | 2.9483467 | 42.4859484 | 1156 | *P. c. asparagoides* |  |  | Collectors VH, JB |
| 21-Ped-32 | Pic_Neolous | 2.9483033 | 42.4859950 | 1153 | *P. c. asparagoides* |  |  | Collectors VH, JB |
| 21-Ped-33 | Pic_Neolous | 2.9448266 | 42.4852601 | 1127 | *P. c. asparagoides* |  |  | Collectors VH, JB |
| 21-Ped-34 | Mijanes | 2.0240900 | 42.7289983 | 1555 | *P. c. comosa* |  |  | Collectors VH,JB,PS |
| 21-Ped-35 | Mijanes | 2.0243084 | 42.7290533 | 1562 | *P. c. comosa* |  |  | Collectors VH,JB,PS |
| 21-Ped-36 | Mijanes | 2.0244434 | 42.7291334 | 1569 | *P. c. comosa* |  |  | Collectors VH,JB,PS |
| 21-Ped-37 | Mijanes | 2.0247084 | 42.7290367 | 1572 | *P. c. comosa* |  |  | Collectors VH,JB,PS |
| 21-Ped-38 | Mijanes | 2.0248251 | 42.7290201 | 1572 | *P. c. comosa* |  |  | Collectors VH,JB,PS |
| 21-Ped-39 | Mijanes | 2.0252534 | 42.7288083 | 1572 | *P. c. comosa* |  |  | Collectors VH,JB,PS |
| 21-Ped-40 | Mijanes | 2.0254817 | 42.7288367 | 1582 | *P. c. comosa* |  |  | Collectors VH,JB,PS |
| 21-Ped-41 | Mijanes | 2.0253850 | 42.7289183 | 1581 | *P. c. comosa* |  |  | Collectors VH,JB,PS |
| 21-Ped-42 | Mijanes | 2.0253134 | 42.7292167 | 1601 | *P. c. comosa* |  |  | Collectors VH,JB,PS |
| 21-Ped-43 | Mijanes | 2.0253650 | 42.7293550 | 1612 | *P. c. comosa* |  |  | Collectors VH,JB,PS |
| 21-Ped-44 | Mijanes | 2.0250583 | 42.7295233 | 1614 | *P. c. comosa* |  |  | Collectors VH,JB,PS |
| 21-Ped-45 | Mijanes | 2.0248966 | 42.7296283 | 1618 | *P. c. comosa* |  |  | Collectors VH,JB,PS |
| 21-Ped-46 | Mijanes | 2.0245867 | 42.7296267 | 1611 | *P. c. comosa* |  |  | Collectors VH,JB,PS |
| 21-Ped-47 | Mijanes | 2.0242583 | 42.7297533 | 1610 | *P. c. comosa* |  |  | Collectors VH,JB,PS |
| 21-Ped-48 | Mijanes | 2.0238917 | 42.7299800 | 1615 | *P. c. comosa* |  |  | Collectors VH,JB,PS |
| 21-Ped-49 | Pic_des_Salines | 2.75129008 | 42.425822 | 1359 | *P. c. asparagoides* |  |  | Problème GPS. Voir C. Hurson |
| 21-Ped-50 | Pic_des_Salines | 2.7512698 | 42.4257481 | 1363 | *P. c. asparagoides* |  |  | Problème GPS. Voir C. Hurson |
| 21-Ped-51 | Pic_des_Salines | 2.751554 | 42.4258129 | 1369 | *P. c. asparagoides* |  |  | Problème GPS. Voir C. Hurson |
| 21-Ped-52 | Pic_des_Salines | 2.75129008 | 42.425822 | 1359 | *P. c. asparagoides* |  |  | Problème GPS. Voir C. Hurson |
| 21-Ped-53 | Pic_des_Salines | 2.7511663 | 42.4256084 | 1370 | *P. c. asparagoides* |  |  | Problème GPS. Voir C. Hurson |
| 21-Ped-54 | Pic_des_Salines | 2.75129008 | 42.425822 | 1359 | *P. c. asparagoides* |  |  | Problème GPS. Voir C. Hurson |
| 21-Ped-55 | Pic_des_Salines | 2.75129008 | 42.425822 | 1359 | *P. c. asparagoides* |  |  | Problème GPS. Voir C. Hurson |
| 21-Ped-56 | Pic_des_Salines | 2.75129008 | 42.425822 | 1359 | *P. c. asparagoides* |  |  | Problème GPS. Voir C. Hurson |
| 21-Ped-57 | Pic_des_Salines | 2.75129008 | 42.425822 | 1359 | *P. c. asparagoides* |  |  | Problème GPS. Voir C. Hurson |
| 21-Ped-58 | Pic_des_Salines | 2.75129008 | 42.425822 | 1359 | *P. c. asparagoides* |  |  | Problème GPS. Voir C. Hurson |
| 21-Ped-59 | Pic_des_Salines | 2.7510827 | 42.4259754 | 1344 | *P. c. asparagoides* |  |  | Problème GPS. Voir C. Hurson |
| 21-Ped-60 | Pic_des_Salines | 2.7513776 | 42.4259652 | 1350 | *P. c. asparagoides* |  |  | Problème GPS. Voir C. Hurson |
| 21-Ped-61 | Pic_des_Salines | 2.75129008 | 42.425822 | 1359 | *P. c. asparagoides* |  |  | Problème GPS. Voir C. Hurson |
| 21-Ped-62 | Pic_des_Salines | 2.75129008 | 42.425822 | 1359 | *P. c. asparagoides* |  |  | Problème GPS. Voir C. Hurson |
| 21-Ped-63 | Pic_des_Salines | 2.75129008 | 42.425822 | 1359 | *P. c. asparagoides* |  |  | Problème GPS. Voir C. Hurson |
| 21-Ped-64 | Pic_des_Salines | 2.75129008 | 42.425822 | 1359 | *P. c. asparagoides* |  |  | Problème GPS. Voir C. Hurson |
| 21-Ped-65 | Py-Mantet | 2.2825650676551 | 42.484016387565 | 1966 | *P. c. comosa* |  |  | Collector V. Hinoux |
| 21-Ped-66 | Py-Mantet | 2.2820542843718 | 42.483995172425 | 1976 | *P. c. comosa* |  |  | Collector V. Hinoux |
| 21-Ped-67 | Py-Mantet | 2.2795075 | 42.4838018 | 2039 | *P. c. comosa* |  |  | Collector V. Hinoux |
| 21-Ped-68 | Py-Mantet | 2.272030017557 | 42.483859795359 | 2041 | *P. c. comosa* |  |  | Collector V. Hinoux |
| 21-Ped-69 | Py-Mantet | 2.2797209179553 | 42.483755362305 | 2033 | *P. c. comosa* |  |  | Collector V. Hinoux |
| 21-Ped-70 | Py-Mantet | 2.2794061430088 | 42.483618293017 | 2026 | *P. c. comosa* |  |  | Collector V. Hinoux |
| 21-Ped-71 | Py-Mantet | 2.279724017007 | 42.483485206668 | 2018 | *P. c. comosa* |  |  | Collector V. Hinoux |
| 21-Ped-72 | Py-Mantet | 2.2788369114772 | 42.483389561172 | 2018 | *P. c. comosa* |  |  | Collector V. Hinoux |
| 21-Ped-73 | Py-Mantet | 2.2788796155902 | 42.482849479962 | 1994 | *P. c. comosa* |  |  | Collector V. Hinoux |
| 21-Ped-74 | Py-Mantet | 2.2785885585551 | 42.482766592906 | 1988 | *P. c. comosa* |  |  | Collector V. Hinoux |
| 21-Ped-75 | Py-Mantet | 2.2776145408408 | 42.482823489259 | 1981 | *P. c. comosa* |  |  | Collector V. Hinoux |
| 21-Ped-76 | Py-Mantet | 2.2773077558001 | 42.481993015331 | 1992 | *P. c. comosa* |  |  | Collector V. Hinoux |
| 21-Ped-77 | Py-Mantet | 2.2779380567093 | 42.483257811312 | 2002 | *P. c. comosa* |  |  | Collector V. Hinoux |
| 21-Ped-78 | Py-Mantet | 2.2784839859101 | 42.483396342325 | 2009 | *P. c. comosa* |  |  | Collector V. Hinoux |
| 21-Ped-79 | Py-Mantet | 2.2786056487122 | 42.483397109356 | 2011 | *P. c. comosa* |  |  | Collector V. Hinoux |
| 21-Ped-80 | Py-Mantet | 2.2787495752686 | 42.483578133346 | 2017 | *P. c. comosa* |  |  | Collector V. Hinoux |
| 21-Ped-81 | Prats-de-Mollo | 2.403177 | 42.452823 | 2044 | *P. c. comosa* |  |  | Collectors PS. CQ. PG, JB (stagiaire) |
| 21-Ped-82 | Prats-de-Mollo | 2.406630 | 42.453988 | 2059 | *P. c. comosa* |  |  | Collectors PS. CQ. PG, JB(stagiaire) |
| 21-Ped-83 | Prats-de-Mollo | 2.406792 | 42.454238 | 2055 | *P. c. comosa* |  |  | Collectors PS. CQ. PG et JB(stagiaire) |
| 21-Ped-84 | Costabonne | 2.3552168128061 | 42.415166373078 | 1982 | *P. c. comosa* |  |  | Collector P. Gaultier |
| 21-Ped-85 | Costabonne | 2.3548740288715 | 42.415407602417 | 1998 | *P. c. comosa* |  |  | Collector P. Gaultier |
| 21-Ped-86 | Costabonne | 2.3545471820576 | 42.415279676331 | 2010 | *P. c. comosa* |  |  | Collector P. Gaultier |
| 21-Ped-87 | Costabonne | 2.3540834160258 | 42.415466185245 | 2029 | *P. c. comosa* |  |  | Collector P. Gaultier |
| 21-Ped-88 | Costabonne | 2.3534387343935 | 42.415516580803 | 2043 | *P. c. comosa* |  |  | Collector P. Gaultier |
| 21-Ped-89 | Costabonne | 2.3526368918533 | 42.415485030627 | 2091 | *P. c. comosa* |  |  | Collector P. Gaultier |
| 21-Ped-90 | Costabonne | 2.3479765280168 | 42.418349489224 | 2245 | *P. c. comosa* |  |  | Collector P. Gaultier |
| 21-Ped-91 | Costabonne | 2.3481802524942 | 42.41862983409 | 2240 | *P. c. comosa* |  |  | Collector P. Gaultier |
| 21-Ped-92 | Asmaris | 2.401465095643 | 42.455800175047 | 2320 | *P. c. comosa* |  |  | Collector P. Gaultier |
| 21-Ped-93 | Asmaris | 2.4019483746766 | 42.456135919899 | 2313 | *P. c. comosa* |  |  | Collector P. Gaultier |
| 21-Ped-94 | Asmaris | 2.4022753596013 | 42.456281722329 | 2308 | *P. c. comosa* |  |  | Collector P. Gaultier |
| 21-Ped-95 | Asmaris | 2.4023993704624 | 42.456030204936 | 2294 | *P. c. comosa* |  |  | Collector P. Gaultier |
| 21-Ped-96 | Asmaris | 2.4062067953174 | 42.454663112979 | 2108 | *P. c. comosa* |  |  | Collector P. Gaultier |
| 21-Ped-97 | Jujols | 2.236975 | 42.619597 | 1604 | *P. c. comosa* |  |  | Collectors PS. CQ et JB |
| 21-Ped-98 | Jujols | 2.237023 | 42.619577 | 1605 | *P. c. comosa* |  |  | Collectors PS. CQ et JB |
| 21-Ped-99 | Jujols | 2.236900 | 42.619422 | 1610 | *P. c. comosa* |  |  | Collectors PS. CQ et JB |
| 21-Ped-100 | Jujols | 2.236743 | 42.616642 | 1668 | *P. c. comosa* |  |  | Collectors PS. CQ et JB |
| 21-Ped-101 | Jujols | 2.234377 | 42.616438 | 1781 | *P. c. comosa* |  |  | Collectors PS. CQ et JB |
| 21-Ped-102 | Jujols | 2.234530 | 42.616382 | 1765 | *P. c. comosa* |  |  | Collectors PS. CQ et JB |
| 21-Ped-103 | Jujols | 2.234828 | 42.616747 | 1770 | *P. c. comosa* |  |  | Collectors PS. CQ et JB |
| 21-Ped-104 | Jujols | 2.235592 | 42.617080 | 1759 | *P. c. comosa* |  |  | Collectors PS. CQ et JB |
| 21-Ped-105 | Jujols | 2.232928 | 42.620060 | 1681 | *P. c. comosa* |  |  | Collectors PS. CQ et JB |
| 21-Ped-106 | Jujols | 2.232913 | 42.629113 | 1681 | *P. c. comosa* |  |  | Collectors PS. CQ et JB |
| 21-Ped-107 | Jujols | 2.232377 | 42.620108 | 1679 | *P. c. comosa* |  |  | Collectors PS. CQ et JB |
| 21-Ped-108 | Jujols | 2.232347 | 42.620107 | 1680 | *P. c. comosa* |  |  | Collectors PS. CQ et JB |
| 21-Ped-109 | Llo | 2.0987 | 42.42819 | 1851 | *P. c. comosa* |  |  | Collector J.-M. Lewin |
| 21-Ped-110 | Llo | 2.09569 | 42.43339 | 1972 | *P. c. comosa* |  |  | Collector J.-M. Lewin |
| 21-Ped-111 | Llo | 2.09172 | 42.42932 | 1671 | *P. c. comosa* |  |  | Collector J.-M. Lewin |
| 21-Ped-112 | Llo | 2.09172 | 42.42932 | 1663 | *P. c. comosa* |  |  | Collector J.-M. Lewin |
| 21-Ped-113 | Llo | 2.08810 | 42.43282 | 1629 | *P. c. comosa* |  |  | Collector J.-M. Lewin |
| 21-Ped-114 | Llo | 2.08743 | 42.43336 | 1619 | *P. c. comosa* |  |  | Collector J.-M. Lewin |
| 21-Ped-115 | Llo | 2.08656 | 42.43407 | 1631 | *P. c. comosa* |  |  | Collector J.-M. Lewin |
| 21-Ped-116 | Llo | 2.08593 | 42.43437 | 1627 | *P. c. comosa* |  |  | Collector J.-M. Lewin |
| 21-Ped-117 | Llo | 2.08554 | 42.43448 | 1614 | *P. c. comosa* |  |  | Collector J.-M. Lewin |
| 21-Ped-118 | Llo | 2.08483 | 42.43484 | 1607 | *P. c. comosa* |  |  | Collector J.-M. Lewin |
| 21-Ped-119 | Llo | 2.07663 | 42.44409 | 1522 | *P. c. comosa* |  |  | Collector J.-M. Lewin |
| 21-Ped-120 | Llo | 2.07142 | 42.44622 | 1487 | *P. c. comosa* |  |  | Collector J.-M. Lewin |
| 21-Ped-121 | Llo | 2.07117 | 42.44637 | 1489 | *P. c. comosa* |  |  | Collector J.-M. Lewin |
| 21-Ped-122 | Llo | 2.06579 | 42.44942 | 1425 | *P. c. comosa* |  |  | Collector J.-M. Lewin |
| 21-Ped-123 | Llo | 2.06334 | 42.45084 | 1404 | *P. c. comosa* |  |  | Collector J.-M. Lewin |
| 21-Ped-124 | Llo | 2.06326 | 42.45068 | 1407 | *P. c. comosa* |  |  | Collector J.-M. Lewin |
| 21-Ped-125 | Mariailles | 2.410365 | 42.493818 | 1766 | *P. c. asparagoides* |  | x | Collector PS. MM. JML et JB |
| 21-Ped-126 | Mariailles | 2.410058 | 42.492972 | 1794 | *P. c. asparagoides* |  | x | Collector PS. MM. JML et JB |
| 21-Ped-127 | Mariailles | 2.410070 | 42.492973 | 1794 | *P. c. asparagoides* |  |  | Collector PS. MM. JML et JB |
| 21-Ped-128 | Mariailles | 2.409742 | 42.493352 | 1809 | *P. c. asparagoides* |  |  | Collector PS. MM. JML et JB |
| 21-Ped-129 | Mariailles | 2.409815 | 42.493317 | 1811 | *P. c. asparagoides* |  |  | Collector PS. MM. JML et JB |
| 21-Ped-130 | Mariailles | 2.409688 | 42.493462 | 1818 | *P. c. asparagoides* |  | x | Collector PS. MM. JML et JB |
| 21-Ped-131 | Mariailles | 2.409255 | 42.492977 | 1837 | *P. c. asparagoides* |  | x | Collector PS. MM. JML et JB |
| 21-Ped-132 | Mariailles | 2.409432 | 42.492798 | 1822 | *P. c. asparagoides* |  |  | Collector PS. MM. JML et JB |
| 21-Ped-133 | Mariailles | 2.409403 | 42.492777 | 1823 | *P. c. asparagoides* |  | x | Collector PS. MM. JML et JB |
| 21-Ped-134 | Mariailles | 2.410170 | 42.492995 | 1784 | *P. c. asparagoides* |  |  | Collector PS. MM. JML et JB |
| 21-Ped-135 | Mariailles | 2.411377 | 42.491872 | 1790 | *P. c. asparagoides* |  |  | Collector PS. MM. JML et JB. Prélevé pour le génome de référence |
| 21-Ped-136 | Mariailles | 2.411452 | 42.491690 | 1792 | *P. c. asparagoides* |  | x | Collector PS. MM. JML et JB |
| 21-Ped-137 | Mariailles | 2.411470 | 42.491765 | 1791 | *P. c. asparagoides* |  | x | Collector PS. MM. JML et JB |
| 21-Ped-138 | Nohèdes | 2.265107 | 42.616470 | 1640 | *P. c. comosa* |  |  | Collector PS. MM. JML et JB |
| 21-Ped-139 | Nohèdes | 2.265107 | 42.616470 | 1640 | *P. c. comosa* |  |  | Collector PS. MM. JML et JB |
| 21-Ped-140 | Nohèdes | 2.265392 | 42.616258 | 1641 | *P. c. comosa* |  |  | Collector PS. MM. JML et JB |
| 21-Ped-141 | Nohèdes | 2.265207 | 42.616452 | 1641 | *P. c. comosa* |  |  | Collector PS. MM. JML et JB |
| 21-Ped-142 | Nohèdes | 2.264968 | 42.616242 | 1650 | *P. c. comosa* |  |  | Collector PS. MM. JML et JB |
| 21-Ped-143 | Nohèdes | 2.264788 | 42.616313 | 1650 | *P. c. comosa* |  |  | Collector PS. MM. JML et JB |
| 21-Ped-144 | Nohèdes | 2.265372 | 42.616558 | 1635 | *P. c. comosa* |  |  | Collector PS. MM. JML et JB |
| 21-Ped-145 | Nohèdes | 2.265578 | 42.616570 | 1629 | *P. c. comosa* |  |  | Collector PS. MM. JML et JB |
| 21-Ped-146 | Capolat | 1.747809458257437 | 42.07796980353451 | NA | *P. c. comosa* |  |  | Collector Pere Aymerich |
| 21-Ped-147 | Capolat | 1.747809458257437 | 42.07796980353451 | NA | *P. c. comosa* |  |  | Collector Pere Aymerich |
| 21-Ped-148 | Capolat | 1.747809458257437 | 42.07796980353451 | NA | *P. c. comosa* |  |  | Collector Pere Aymerich |
| 21-Ped-149 | Capolat | 1.747809458257437 | 42.07796980353451 | NA | *P. c. comosa* |  |  | Collector Pere Aymerich |
| 21-Ped-150 | Capolat | 1.747809458257437 | 42.07796980353451 | NA | *P. c. comosa* |  |  | Collector Pere Aymerich |
| 21-Ped-151 | Capolat | 1.747809458257437 | 42.07796980353451 | NA | *P. c. comosa* |  |  | Collector Pere Aymerich |
| 21-Ped-152 | Capolat | 1.747809458257437 | 42.07796980353451 | NA | *P. c. comosa* |  |  | Collector Pere Aymerich |
| 21-Ped-153 | Capolat | 1.747809458257437 | 42.07796980353451 | NA | *P. c. comosa* |  |  | Collector Pere Aymerich |
| 21-Ped-154 | Capolat | 1.747809458257437 | 42.07796980353451 | NA | *P. c. comosa* |  |  | Collector Pere Aymerich |
| 21-Ped-155 | Capolat | 1.747809458257437 | 42.07796980353451 | NA | *P. c. comosa* |  |  | Collector Pere Aymerich |
| 21-Ped-156 | Capolat | 1.747809458257437 | 42.07796980353451 | NA | *P. c. comosa* |  |  | Collector Pere Aymerich |
| 21-Ped-157 | Capolat | 1.747809458257437 | 42.07796980353451 | NA | *P. c. comosa* |  |  | Collector Pere Aymerich |
| 21-Ped-158 | Capolat | 1.747809458257437 | 42.07796980353451 | NA | *P. c. comosa* |  |  | Collector Pere Aymerich |
| 23-Ped-159 | Pic_des_Salines | 2.75129008 | 42.425822 | 1359 | *P. c. asparagoides* | x | x | Collector VH, JB, PS |
| 23-Ped-160 | Pic_des_Salines | 2.75129008 | 42.425822 | 1359 | *P. c. asparagoides* | x | x | Collector VH, JB, PS |
| 23-Ped-161 | Pic_des_Salines | 2.75129008 | 42.425822 | 1359 | *P. c. asparagoides* | x | x | Collector VH, JB, PS |
| 23-Ped-162 | Pic_des_Salines | 2.75129008 | 42.425822 | 1359 | *P. c. asparagoides* |  |  | Collector VH, JB, PS |
| 23-Ped-163 | Pic_des_Salines | 2.75129008 | 42.425822 | 1359 | *P. c. asparagoides* |  |  | Collector VH, JB, PS |
| 23-Ped-164 | Pic_des_Salines | 2.75129008 | 42.425822 | 1359 | *P. c. asparagoides* |  |  | Collector VH, JB, PS |
| 23-Ped-165 | Pic_des_Salines | 2.75129008 | 42.425822 | 1359 | *P. c. asparagoides* |  |  | Collector VH, JB, PS |
| 23-Ped-166 | Pic_des_Salines | 2.75129008 | 42.425822 | 1359 | *P. c. asparagoides* |  |  | Collector VH, JB, PS |
| 23-Ped-167 | Mijanes | 2.024791027 | 42.72925923 | 1590 | *P. c. comosa* | x | x | Collector JB,  PS |
| 23-Ped-168 | Mijanes | 2.024791027 | 42.72925923 | 1590 | *P. c. comosa* | x | x | Collector JB,  PS |
| 23-Ped-169 | Mijanes | 2.024791027 | 42.72925923 | 1590 | *P. c. comosa* | x | x | Collector JB,  PS |
| 23-Ped-170 | Mijanes | 2.024791027 | 42.72925923 | 1590 | *P. c. comosa* |  | x | Collector JB,  PS |
| 23-Ped-171 | Mijanes | 2.024791027 | 42.72925923 | 1590 | *P. c. comosa* |  | x | Collector JB,  PS |
| 23-Ped-172 | Mijanes | 2.024791027 | 42.72925923 | 1590 | *P. c. comosa* |  | x | Collector JB,  PS |
| 23-Ped-173 | Mijanes | 2.024791027 | 42.72925923 | 1590 | *P. c. comosa* |  | x | Collector JB,  PS |
| 23-Ped-174 | Mijanes | 2.024791027 | 42.72925923 | 1590 | *P. c. comosa* |  | x | Collector JB,  PS |
| 23-Ped-175 | Pic_Neolous | 2.946987606 | 42.48383476 | 1205 | *P. c. asparagoides* | x | x | Collector VH, AG, JAM, PS |
| 23-Ped-176 | Pic_Neolous | 2.946987606 | 42.48383476 | 1205 | *P. c. asparagoides* | x | x | Collector VH, AG, JAM, PS |
| 23-Ped-177 | Pic_Neolous | 2.946987606 | 42.48383476 | 1205 | *P. c. asparagoides* | x | x | Collector VH, AG, JAM, PS |
| 23-Ped-178 | Pic_Neolous | 2.946987606 | 42.48383476 | 1205 | *P. c. asparagoides* |  | x | Collector VH, AG, JAM, PS |
| 23-Ped-179 | Pic_Neolous | 2.946987606 | 42.48383476 | 1205 | *P. c. asparagoides* |  | x | Collector VH, AG, JAM, PS |
| 23-Ped-180 | Pic_Neolous | 2.946987606 | 42.48383476 | 1205 | *P. c. asparagoides* |  | x | Collector VH, AG, JAM, PS |
| 23-Ped-181 | Pic_Neolous | 2.946987606 | 42.48383476 | 1205 | *P. c. asparagoides* |  | x | Collector VH, AG, JAM, PS |
| 23-Ped-182 | Pic_Neolous | 2.946987606 | 42.48383476 | 1205 | *P. c. asparagoides* |  | x | Collector VH, AG, JAM, PS |
| 23-Ped-183 | Py-Mantet | 2.278750787 | 42.48337397 | 2007 | *P. c. comosa* | x | x | Collector LJ, JB, PS |
| 23-Ped-184 | Py-Mantet | 2.278750787 | 42.48337397 | 2007 | *P. c. comosa* |  | x | Collector LJ, JB, PS |
| 23-Ped-185 | Py-Mantet | 2.278750787 | 42.48337397 | 2007 | *P. c. comosa* | x | x | Collector LJ, JB, PS |
| 23-Ped-186 | Py-Mantet | 2.278750787 | 42.48337397 | 2007 | *P. c. comosa* | x | x | Collector LJ, JB, PS |
| 23-Ped-187 | Py-Mantet | 2.278750787 | 42.48337397 | 2007 | *P. c. comosa* |  | x | Collector LJ, JB, PS |
| 23-Ped-188 | Py-Mantet | 2.278750787 | 42.48337397 | 2007 | *P. c. comosa* |  | x | Collector LJ, JB, PS |
| 23-Ped-189 | Py-Mantet | 2.278750787 | 42.48337397 | 2007 | *P. c. comosa* |  | x | Collector LJ, JB, PS |
| 23-Ped-190 | Py-Mantet | 2.278750787 | 42.48337397 | 2007 | *P. c. comosa* |  | x | Collector LJ, JB, PS |
| 23-Ped-191 | Llo | 2.081739375 | 42.43823625 | 1601 | *P. c. comosa* | x | x | Collector JML, JB, PS |
| 23-Ped-192 | Llo | 2.081739375 | 42.43823625 | 1601 | *P. c. comosa* | x | x | Collector JML, JB, PS |
| 23-Ped-193 | Llo | 2.081739375 | 42.43823625 | 1601 | *P. c. comosa* |  | x | Collector JML, JB, PS |
| 23-Ped-194 | Llo | 2.081739375 | 42.43823625 | 1601 | *P. c. comosa* |  | x | Collector JML, JB, PS |
| 23-Ped-195 | Llo | 2.081739375 | 42.43823625 | 1601 | *P. c. comosa* |  | x | Collector JML, JB, PS |
| 23-Ped-196 | Llo | 2.081739375 | 42.43823625 | 1601 | *P. c. comosa* |  | x | Collector JML, JB, PS |
| 23-Ped-197 | Llo | 2.081739375 | 42.43823625 | 1601 | *P. c. comosa* |  | x | Collector JML, JB, PS |
| 23-Ped-198 | Llo | 2.081739375 | 42.43823625 | 1601 | *P. c. comosa* | x | x | Collector JML, JB, PS |
| 23-Ped-199 | Mariailles | 2.410176692 | 42.49282831 | 1802 | *P. c. asparagoides* | x | x | Collector JML, JB, PS |
| 23-Ped-200 | Mariailles | 2.410176692 | 42.49282831 | 1802 | *P. c. asparagoides* | x | x | Collector JML, JB, PS |

| A-Fst/Distance | East Albères | West Albères | Mijanès | Pic des Salines | Py-Mantet | Prats-de-Mollo | Jujols | Llo | Mariailles | Nohèdes | Capolat |
| --- | --- | --- | --- | --- | --- | --- | --- | --- | --- | --- | --- |
| East Albères |  | 6 | 86 | 23 | 61 | 53 | 66 | 77 | 50 | 64 | 113 |
| West Albères | 0.062 |  | 80 | 17 | 55 | 47 | 60 | 71 | 44 | 58 | 108 |
| Mijanès | 0.246 | 0.28 |  | 68 | 34 | 44 | 21 | 33 | 41 | 23 | 76 |
| Pic des Salines | 0.235 | 0.262 | 0.393 |  | 39 | 31 | 47 | 55 | 29 | 45 | 91 |
| Py-Mantet | 0.242 | 0.26 | 0.376 | 0.295 |  | 10 | 16 | 17 | 11 | 15 | 63 |
| Prats-de-Mollo | 0.197 | 0.217 | 0.343 | 0.241 | 0.084 |  | 24 | 24 | 7 | 22 | 65 |
| Jujols | 0.178 | 0.218 | 0.189 | 0.328 | 0.31 | 0.272 |  | 24 | 20 | 3 | 72 |
| Llo | 0.227 | 0.244 | 0.364 | 0.289 | 0.121 | 0.098 | 0.288 |  | 28 | 25 | 49 |
| Mariailles | 0.194 | 0.214 | 0.345 | 0.247 | 0.142 | 0.072 | 0.27 | 0.136 |  | 18 | 71 |
| Nohèdes | 0.197 | 0.241 | 0.209 | 0.361 | 0.333 | 0.29 | 0.065 | 0.313 | 0.293 |  | 73 |
| Capolat | 0.371 | 0.4 | 0.477 | 0.48 | 0.381 | 0.345 | 0.423 | 0.362 | 0.376 | 0.466 |  |

**Supplementary Table S5:** Pairwise genetic differentiation (**A-***F*ST values, **B-***G*’ST; Meirmans & Hedrick, 2011) and pairwise geographic distances (in km). All values were found to be statistically significant (*p* < 0.05).

| B-G'st(Nei)/Distance | East Albères | West Albères | Mijanès | Pic des Salines | Py-Mantet | Prats-de-Mollo | Jujols | Llo | Mariailles | Nohèdes | Capolat |
| --- | --- | --- | --- | --- | --- | --- | --- | --- | --- | --- | --- |
| East Albères |  | 6 | 86 | 23 | 61 | 53 | 66 | 77 | 50 | 64 | 113 |
| West Albères | 0.061 |  | 80 | 17 | 55 | 47 | 60 | 71 | 44 | 58 | 108 |
| Mijanès | 0.245 | 0.278 |  | 68 | 34 | 44 | 21 | 33 | 41 | 23 | 76 |
| Pic des Salines | 0.235 | 0.264 | 0.392 |  | 39 | 31 | 47 | 55 | 29 | 45 | 91 |
| Py-Mantet | 0.24 | 0.259 | 0.372 | 0.294 |  | 10 | 16 | 17 | 11 | 15 | 63 |
| Prats-de-Mollo | 0.196 | 0.216 | 0.34 | 0.241 | 0.083 |  | 24 | 24 | 7 | 22 | 65 |
| Jujols | 0.177 | 0.214 | 0.187 | 0.322 | 0.303 | 0.267 |  | 24 | 20 | 3 | 72 |
| Llo | 0.225 | 0.243 | 0.361 | 0.288 | 0.121 | 0.098 | 0.282 |  | 28 | 25 | 49 |
| Mariailles | 0.194 | 0.214 | 0.344 | 0.245 | 0.14 | 0.072 | 0.267 | 0.135 |  | 18 | 71 |
| Nohèdes | 0.199 | 0.239 | 0.208 | 0.347 | 0.323 | 0.284 | 0.066 | 0.304 | 0.288 |  | 73 |
| Capolat | 0.374 | 0.409 | 0.478 | 0.478 | 0.384 | 0.348 | 0.416 | 0.365 | 0.373 | 0.443 |  |

**Supplementary Table S6:** Summary of the phenotypic and genetic clines

|  | Magenta Score Cline | Yellow Score Cline | SNMF-K=4-light blue cluster membership | ADMIXTURE-K=4-pink cluster membership |
| --- | --- | --- | --- | --- |
| Center | 58.94 | 49.55 | 82.11 | 79.82 |
| Width | 4.41 | 7.37 | 43.68 | 33.18 |

**Supplementary Table S7:** Summary of the selected clines with a delta of frequency above 0.6 between the Westernmost (11-Capolat) and the Easternmost (1-East Albères) populations, a center between 56 and 70 km, and a width lower or equal to 33 km.

| snp_id | center | width | model | chromosome | basepair |
| --- | --- | --- | --- | --- | --- |
| X83646_136 | 96.2854714799562 | 9.46043155154884 | 2 | 1 | 68762755 |
| X150450_174 | 58.9430805288003 | 22.3590599386019 | 1 | 1 | 120545094 |
| X150450_315 | 58.2827329049472 | 21.291834207889 | 1 | 1 | 120544953 |
| X2808390_41 | 78.7748820945439 | 1.28577815141293 | 2 | 100003 | 27346445 |
| X2854310_91 | 72.0899238118015 | 2.78887155263512 | 2 | 100003 | 64080461 |
| X2873861_45 | 66.028200099959 | 28.028553830199 | 1 | 100003 | 78722953 |
| X2873861_47 | 63.3977816406285 | 30.2905933656521 | 1 | 100003 | 78722955 |
| X2873861_69 | 66.1920848714346 | 28.6361152302212 | 1 | 100003 | 78722977 |
| X2873861_70 | 65.3229623132906 | 24.5176292827281 | 1 | 100003 | 78722978 |
| X2873861_161 | 65.6842110629788 | 24.9124419040063 | 1 | 100003 | 78723069 |
| X2873861_195 | 63.435828535224 | 23.2288479762671 | 2 | 100003 | 78723103 |
| X2888542_124 | 61.203438288427 | 3.1273284756523 | 2 | 100003 | 90567826 |
| X2915878_250 | 60.306959313203 | 1.42615714854433 | 1 | 100003 | 111479413 |
| X2954665_310 | 57.5196901120227 | 24.5156254105812 | 1 | 100003 | 138885262 |
| X3019355_37 | 56.5525433978332 | 2.3499690912931 | 2 | 100010 | 21994304 |
| X3028027_78 | 69.9003887035511 | 6.80817325208366 | 2 | 100010 | 29251601 |
| X3028265_174 | 71.8328407738076 | 2.89217921117123 | 2 | 100010 | 29418947 |
| X3043844_207 | 56.991472636135 | 2.54392065897513 | 2 | 100010 | 42435558 |
| X3054928_150 | 73.2866579114023 | 0.0542292823734396 | 2 | 100010 | 51722519 |
| X3087051_108 | 59.503569423749 | 3.30772780352999 | 2 | 100010 | 76227020 |
| X3094542_261 | 56.2481459748414 | 1.55569134373919 | 2 | 100010 | 82159104 |
| X3115443_375 | 56.6947734966267 | 7.57917432607829 | 2 | 100010 | 97278951 |
| X3116203_224 | 64.2847652579148 | 7.29547092571208 | 2 | 100010 | 97795550 |
| X3163233_379 | 65.3680639724587 | 6.21239324224399 | 2 | 100020 | 22814671 |
| X3251300_193 | 56.5344190261957 | 22.9586861929045 | 1 | 100028 | 26128527 |
| X3256577_152 | 63.2170021604616 | 0.618054530377458 | 1 | 100028 | 28998033 |
| X1875852_459 | 68.6244042067336 | 1.5054770326037 | 2 | 1188 | 82881627 |
| X2523574_25 | 58.9962717014359 | 20.1742909415588 | 2 | 15494 | 4935250 |
| X2523598_239 | 56.365546029497 | 0.92483567905327 | 2 | 15494 | 4938401 |
| X2533744_74 | 57.216654199301 | 4.09665218202187 | 2 | 15494 | 12668034 |
| X2533744_180 | 58.156141062531 | 0.155524008527884 | 1 | 15494 | 12668140 |
| X2559607_281 | 56.1705365553848 | 2.0052299165678 | 2 | 15497 | 17191730 |
| X2587438_77 | 75.4488130384825 | 0.485448265992772 | 2 | 15662 | 8999755 |
| X2688285_209 | 84.5443131875748 | 1.69536071284909 | 2 | 15662 | 88343623 |
| X2715809_101 | 79.1878011648103 | 2.419772549813 | 2 | 15662 | 109734624 |
| X372312_126 | 81.218230008563 | 0.217321470777681 | 1 | 157 | 24069337 |
| X399271_129 | 56.7641908081833 | 0.0416948811326532 | 1 | 252 | 55396 |
| X418051_87 | 66.8682550206757 | 5.66108684225396 | 2 | 252 | 113392 |
| X422401_64 | 65.6402148436294 | 1.61073407542078 | 2 | 252 | 124325 |
| X428184_68 | 70.2843653063895 | 3.70217201610342 | 2 | 252 | 142517 |
| X443654_30 | 77.9958182155385 | 1.00357891836566 | 2 | 252 | 192726 |
| X480031_38 | 61.6806966134342 | 19.1920660725399 | 1 | 439 | 5441596 |
| X508874_331 | 65.2914177292343 | 17.6138755632969 | 2 | 439 | 26042067 |
| X511642_119 | 67.3796287219464 | 31.4039173624171 | 1 | 439 | 28184150 |
| X511642_125 | 56.8303180828982 | 13.8710706831264 | 2 | 439 | 28184156 |
| X527116_21 | 62.5217963134125 | 11.4151289204268 | 1 | 439 | 40371841 |
| X527116_56 | 62.4852775796459 | 12.0704600489998 | 1 | 439 | 40371806 |
| X527116_73 | 59.9191264968574 | 1.66727464116101 | 1 | 439 | 40371789 |
| X527116_84 | 59.736681456557 | 1.10568877934665 | 1 | 439 | 40371778 |
| X527116_156 | 61.129617698698 | 9.44152396936088 | 1 | 439 | 40371706 |
| X527116_180 | 62.5791555569755 | 12.226041480309 | 1 | 439 | 40371682 |
| X527116_266 | 59.7352199125776 | 1.17938120333154 | 1 | 439 | 40371596 |
| X534157_116 | 64.6669487817689 | 8.96229789145318 | 2 | 439 | 44634958 |
| X549556_114 | 64.3162897699317 | 2.26901900844509 | 2 | 439 | 54884685 |
| X625181_195 | 62.0053018850447 | 3.33552227139312 | 2 | 439 | 109135440 |
| X627799_17 | 12.3807763793365 | 5.09757066638575 | 2 | 439 | 111125002 |
| X635608_15 | 81.6853413227483 | 2.51932335270591 | 2 | 439 | 117448245 |
| X637875_351 | 67.6280363617173 | 20.627072184653 | 2 | 439 | 118842244 |
| X713965_414 | 56.4868625586407 | 0.240533836802287 | 2 | 439 | 175329591 |
| X720955_295 | 51.0525271545867 | 0.0734089884616356 | 2 | 439 | 181326196 |
| X838079_284 | 61.6729947726807 | 4.27125929354533 | 2 | 489 | 61872137 |
| X850780_38 | 62.1908556680545 | 9.06626320682083 | 1 | 489 | 70848126 |
| X850780_275 | 62.7701785227837 | 10.3253933107862 | 1 | 489 | 70847889 |
| X871721_191 | 63.2862364514592 | 22.7484925191082 | 2 | 489 | 84768785 |
| X904623_95 | 60.292315497336 | 2.16460046015737 | 2 | 489 | 109432943 |
| X919609_237 | 64.0451772221077 | 32.7406637491281 | 1 | 489 | 120458037 |
| X950026_148 | 59.6741758715122 | 2.20617315269521 | 2 | 489 | 142933842 |
| X954801_28 | 55.707203671051 | 3.85890882343051 | 2 | 489 | 148269477 |
| X997545_153 | 61.3208680880116 | 5.0077875962941 | 2 | 491 | 18561992 |
| X1034271_52 | 58.2069183381077 | 5.48864336781804 | 2 | 499 | 7306036 |
| X1034271_112 | 58.541071076271 | 3.98835216421709 | 2 | 499 | 7306096 |
| X1063887_14 | 61.5548847735803 | 4.62995870214713 | 2 | 499 | 26505394 |
| X1063887_16 | 84.2694584485994 | 0.180320434048946 | 2 | 499 | 26505392 |
| X1063887_36 | 61.4930396230319 | 3.75501966094106 | 2 | 499 | 26505372 |
| X1063887_136 | 60.503574536429 | 1.7644247073072 | 2 | 499 | 26505272 |
| X1063887_156 | 60.5759693402323 | 2.72986749943709 | 2 | 499 | 26505252 |
| X1124661_312 | 59.1224698085608 | 5.54253072359092 | 2 | 545 | 20839292 |
| X1154222_8 | 58.2559987246967 | 3.69878460207344 | 1 | 545 | 46428648 |
| X1229317_244 | 72.5869921705033 | 4.1965162266111 | 2 | 561 | 62645847 |
| X1287034_24 | 59.5676762057932 | 3.2481259178419 | 2 | 561 | 105171109 |
| X1331174_7 | 56.5496573644966 | 2.56757849899134 | 2 | 561 | 136180872 |
| X1345777_103 | 60.0261165550228 | 8.41886819627272 | 2 | 561 | 147785073 |
| X1350852_7 | 66.6541036025773 | 31.8345880986352 | 1 | 561 | 150972532 |
| X1350852_28 | 66.8296367049884 | 31.6325303583265 | 1 | 561 | 150972553 |
| X1350852_51 | 67.0817957616667 | 32.4772743144141 | 1 | 561 | 150972576 |
| X1360306_255 | 58.005329359356 | 9.43179751204509 | 2 | 561 | 157006898 |
| X1465823_57 | 56.4952068653603 | 0.852792828963572 | 1 | 659 | 73492899 |
| X1537117_250 | 57.1332616578214 | 6.55343993254277 | 2 | 659 | 128871315 |
| X1539078_25 | 67.337656590741 | 5.54087431453316 | 2 | 659 | 130402255 |
| X1539078_166 | 72.3231452823818 | 0.862610604004082 | 2 | 659 | 130402396 |
| X1539078_200 | 66.4287044172745 | 6.37092447587919 | 2 | 659 | 130402430 |
| X228700_328 | 65.5773326184581 | 18.3256952950924 | 2 | 66 | 8421984 |
| X228700_330 | 65.0229011353233 | 19.9345158597237 | 2 | 66 | 8421986 |
| X287309_248 | 67.442822594509 | 0.118408876029271 | 2 | 66 | 54835852 |
| X287354_377 | 76.2685890817844 | 1.43240024607824 | 2 | 66 | 54837608 |
| X1991285_352 | 62.7645185829606 | 7.07108726147265 | 2 | 7757 | 2288811 |
| X2014531_417 | 57.0536661161822 | 1.91748941393341 | 2 | 7757 | 25932086 |
| X2025463_285 | 58.1041824015326 | 11.3482945452345 | 2 | 7757 | 33919803 |
| X2152458_132 | 57.3665117854322 | 6.03516349347098 | 1 | 7822 | 4989170 |
| X2152458_245 | 57.1205581468654 | 7.10775853072963 | 1 | 7822 | 4989283 |
| X2152458_289 | 57.0788229538737 | 7.7361179849747 | 1 | 7822 | 4989327 |
| X2152499_272 | 65.7899686879655 | 3.5558206753112 | 2 | 7822 | 4991553 |
| X2189507_306 | 71.6295122538618 | 1.21886419778285 | 2 | 7825 | 902567 |
| X2253386_115 | 55.7443284575664 | 7.55473298091979 | 2 | 7825 | 44086200 |
| X2253386_124 | 56.1011474348549 | 7.13799107337997 | 2 | 7825 | 44086209 |
| X2425388_94 | 61.802668812398 | 23.5596354145941 | 1 | 7825 | 170291312 |
| X2433147_232 | 66.463757137667 | 12.3113599400297 | 2 | 7825 | 176269716 |
| X2433147_299 | 57.5644422656999 | 5.37578434630938 | 2 | 7825 | 176269783 |
| X2438825_38 | 56.3951006020225 | 2.09695425030597 | 2 | 7825 | 181753971 |
| X2448116_360 | 82.6795496848135 | 1.03370036020661 | 2 | 7825 | 189622984 |
| X2501571_212 | 52.3280392613755 | 116.478105076727 | 1 | 7825 | 232395639 |
| X2501980_176 | 59.54833836351 | 1.33628301290805 | 2 | 7825 | 232657273 |
| X2501980_204 | 59.5029937262024 | 1.56313516358769 | 2 | 7825 | 232657245 |
| X1658180_88 | 59.2730333787016 | 2.85671283428506 | 2 | 790 | 34452239 |
| X1658180_219 | 58.9843617230133 | 2.01842413905238 | 2 | 790 | 34452370 |
| X1658180_282 | 59.2360413497478 | 2.88232833153602 | 2 | 790 | 34452433 |
| X1658180_392 | 59.0415180284948 | 1.56859315445574 | 2 | 790 | 34452543 |
| X321077_74 | 55.8459573962157 | 29.8715876247255 | 2 | 89 | 20116217 |

**Supplementary Table S8**: Mapping percentage of the RNA-seq sequences against the reference genome

| Individual_ID | Mapping percentage |
| --- | --- |
| 23-Ped-159 | 78.74 |
| 23-Ped-160 | 68.67 |
| 23-Ped-161 | 89.73 |
| 23-Ped-167 | 87.31 |
| 23-Ped-168 | 90.28 |
| 23-Ped-169 | 88.26 |
| 23-Ped-175 | 89.7 |
| 23-Ped-176 | 87.68 |
| 23-Ped-177 | 89.11 |
| 23-Ped-183 | 87.08 |
| 23-Ped-185 | 88.76 |
| 23-Ped-186 | 86.34 |
| 23-Ped-191 | 88.74 |
| 23-Ped-192 | 86.31 |
| 23-Ped-198 | 90.31 |
| 23-Ped-199 | 87.74 |
| 23-Ped-200 | 89.69 |
| 23-Ped-201 | 87.68 |
| 23-Ped-202 | 78.61 |
| 23-Ped-203 | 64.58 |
| Mean | 85.266 |

**Supplementary Figures**

**Supplementary Figure S1: Genome size estimation**

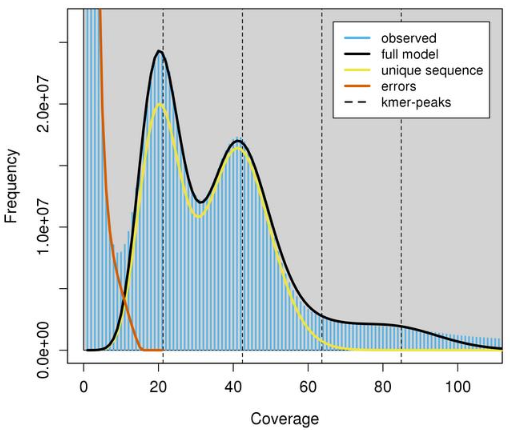

**Supplementary Figure S2: A)**Values of the cross-entropy criterion for a number of clusters ranging from *K*=1 to *K*=12. The optimal number of *K* with Admixture was found to be 5. **B)** Values of the cross-entropy criterion for a number of clusters ranging from *K*=1 to *K*=12. The optimal number of *K* with SNMF was found to be 6. **C)** Barplot displaying individual ancestry coefficients obtained from ADMIXTURE for 156 individuals and 28,271 SNPs for *K*=2 to *K*=7. The optimal value was found to be 5.

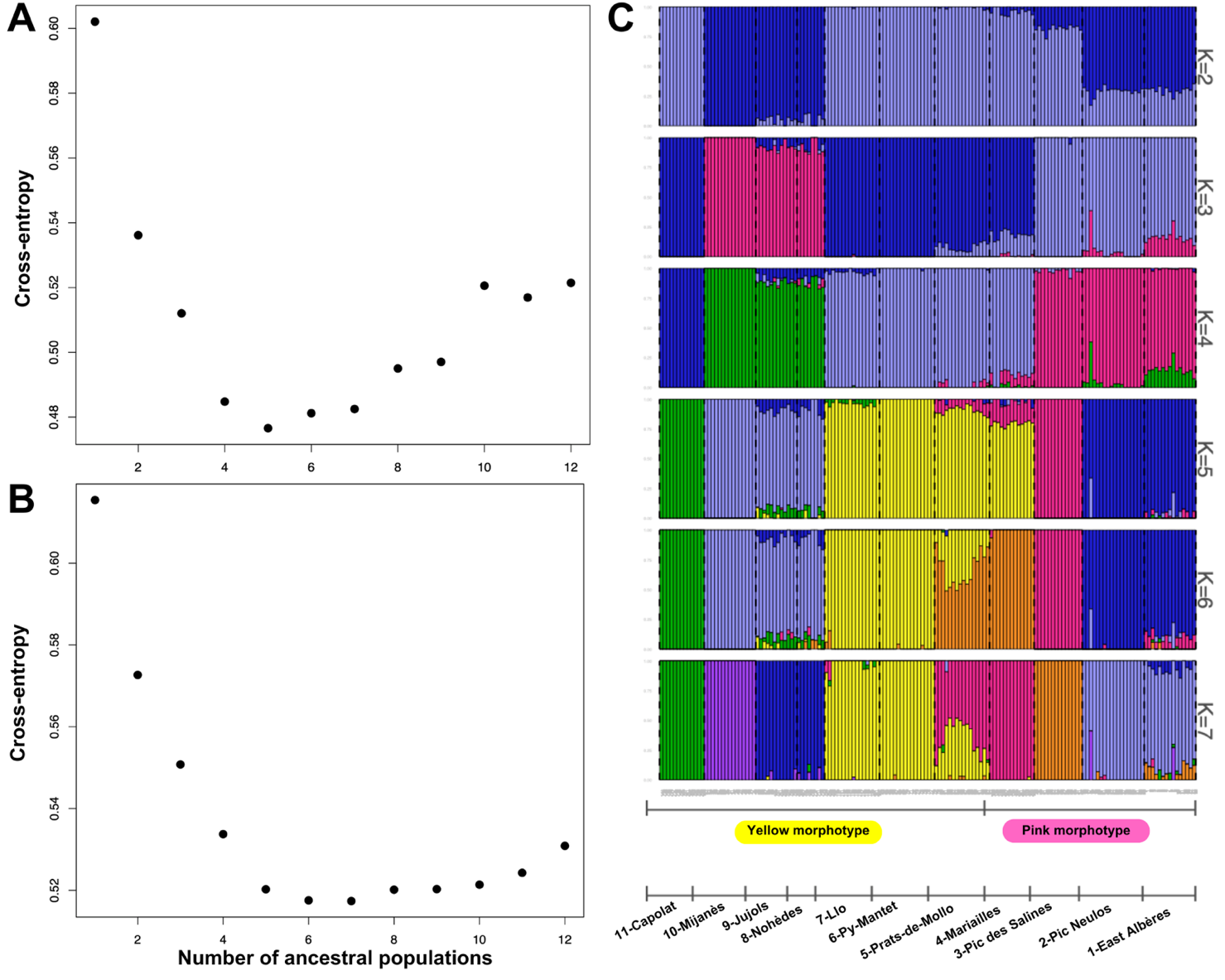

**Supplementary Figure S3:** Co-ancestry matrix between each pair of individuals inferred using fineRADstructure. Each pixel depicts the magnitude of the individual co-ancestry coefficient between two individuals. Low co-ancestry coefficients values are depicted by yellow colors, whereas high values are indicated by darker colors.

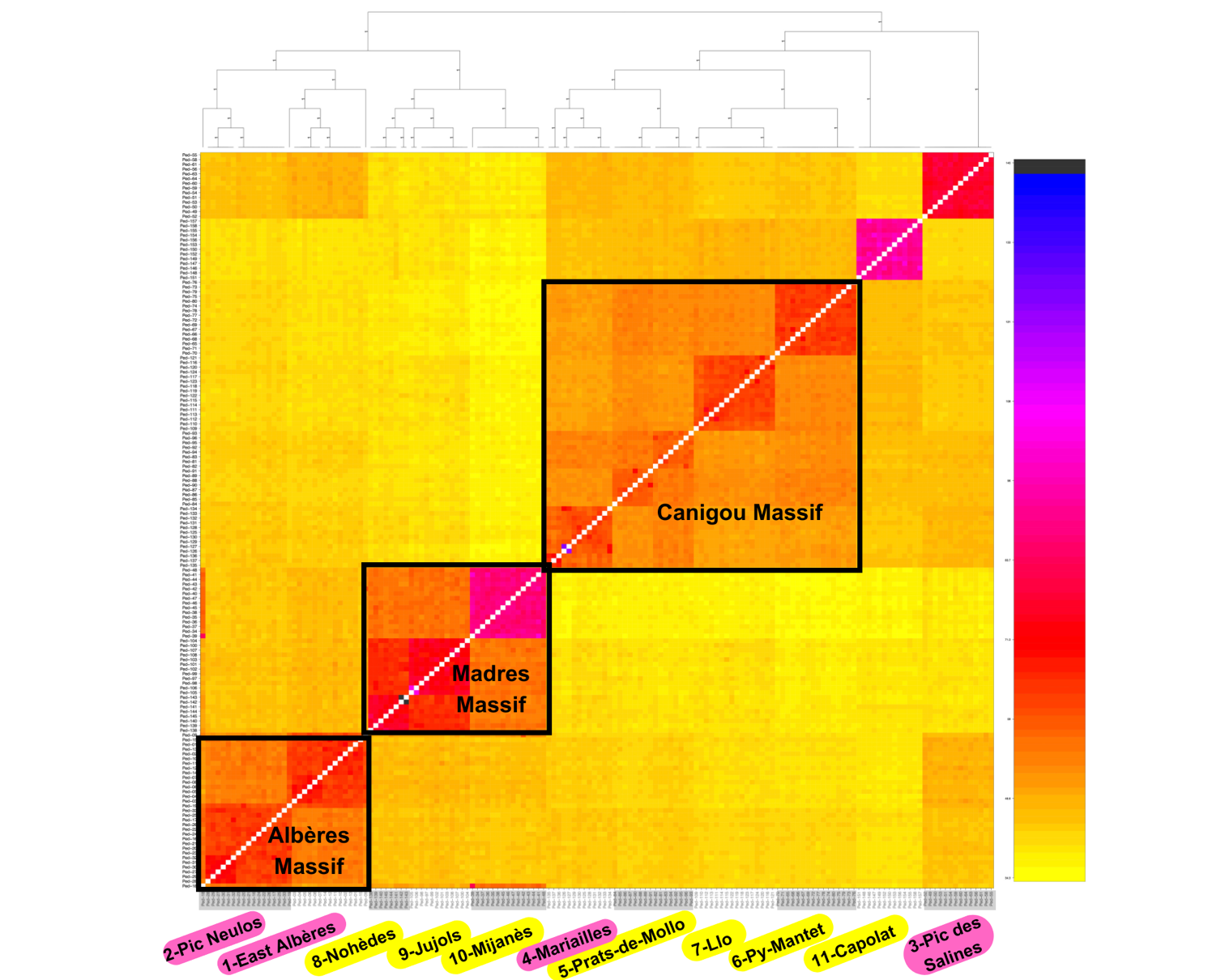

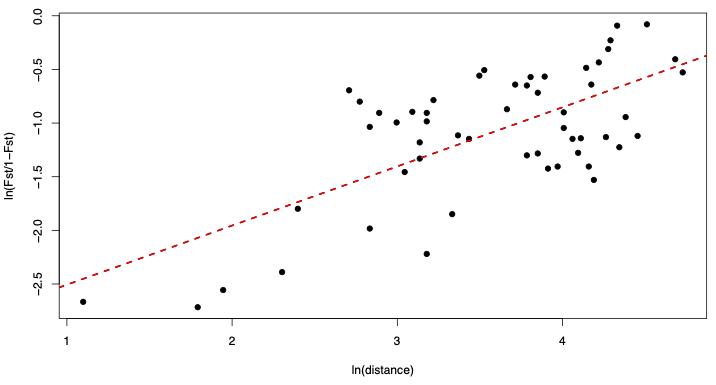
**Supplementary Figure S4:** Pattern of Isolation By Distance (IBD) across the data set. Linearized *F*_ST_ values (i.e., ln(*F*_ST_/1-*F*_ST_)) are plotted against ln-geographical distances. *Mantel statistic*=0.5167 *p-value*=0.015

**Supplementary Figure S5:** Analysis of introgression with *F*-branch. Values of f_b_ refer to excess allele sharing between the branch *b* on the y-axis and populations on the x-axis.

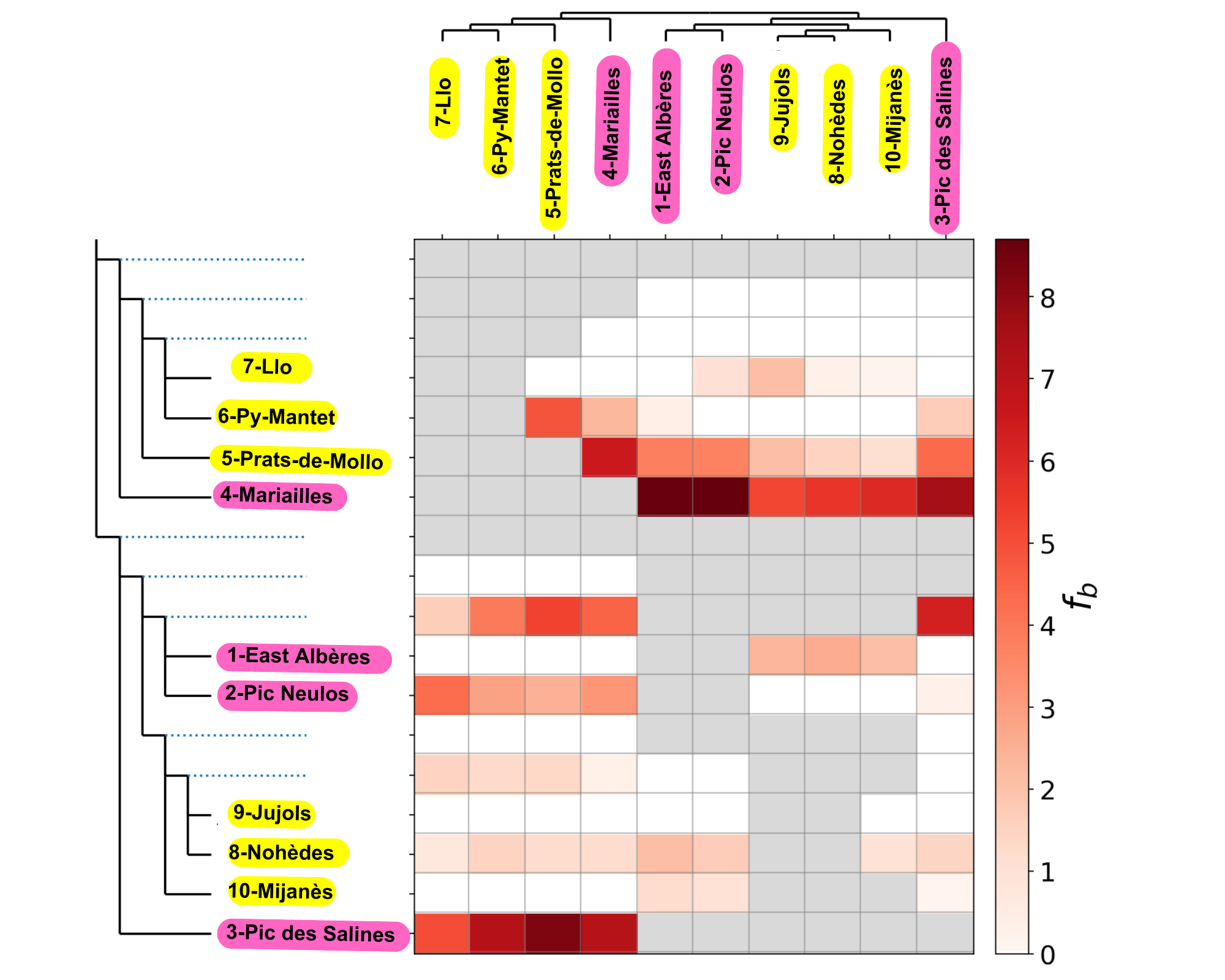

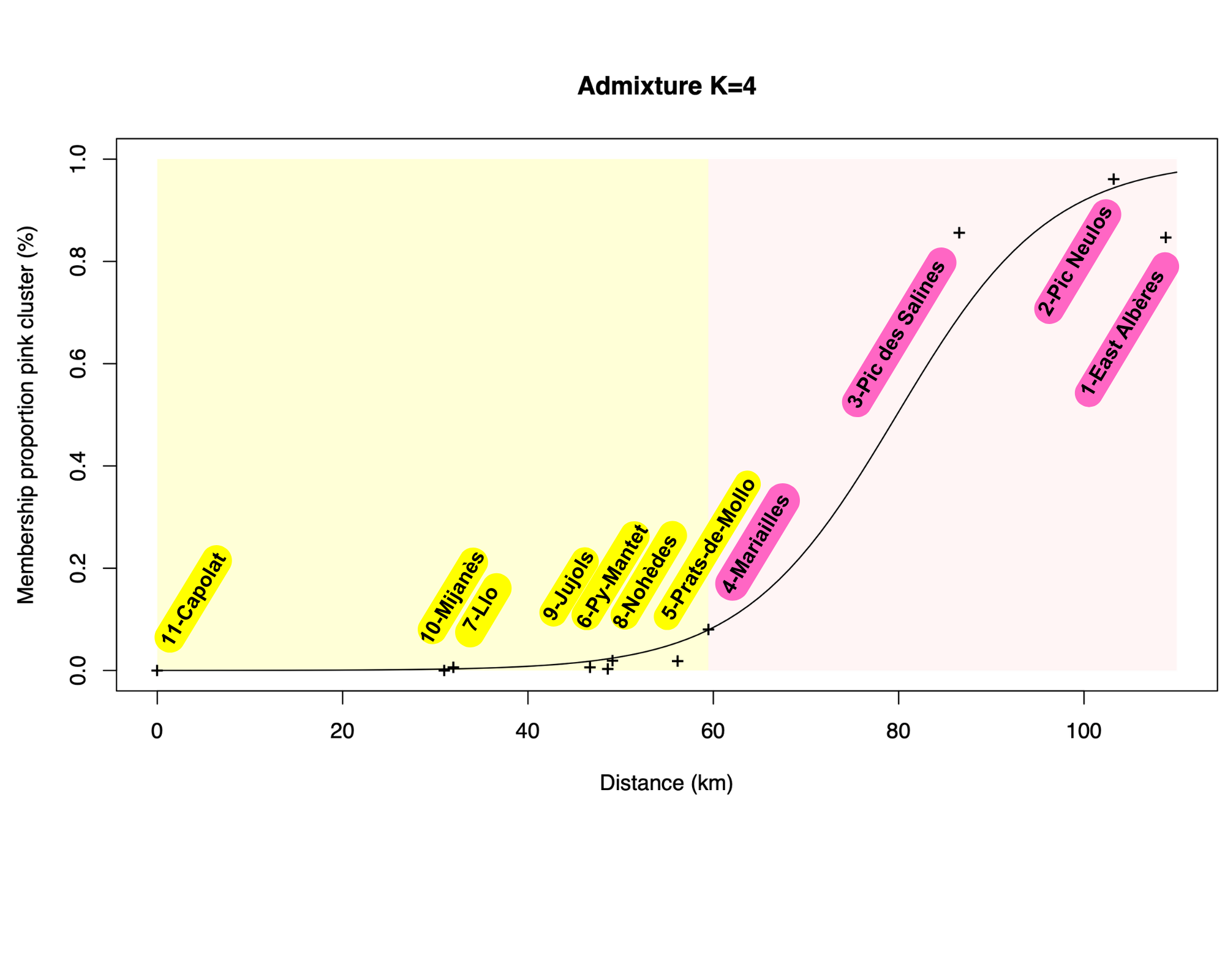
**Supplementary Figure S6:** Genetic cline fitted with HZAR on the membership proportion of the pink cluster displayed by Admixture with *K*=4.

**Supplementary Figure S7:** **A)** Distribution of the values of the center of the cline for all the clines fitted with HZAR on the 28,271 SNPs. The mean of the center is 55.01km. **B)** Distribution of the values of the width of the cline. The mean of the width is 29.60km

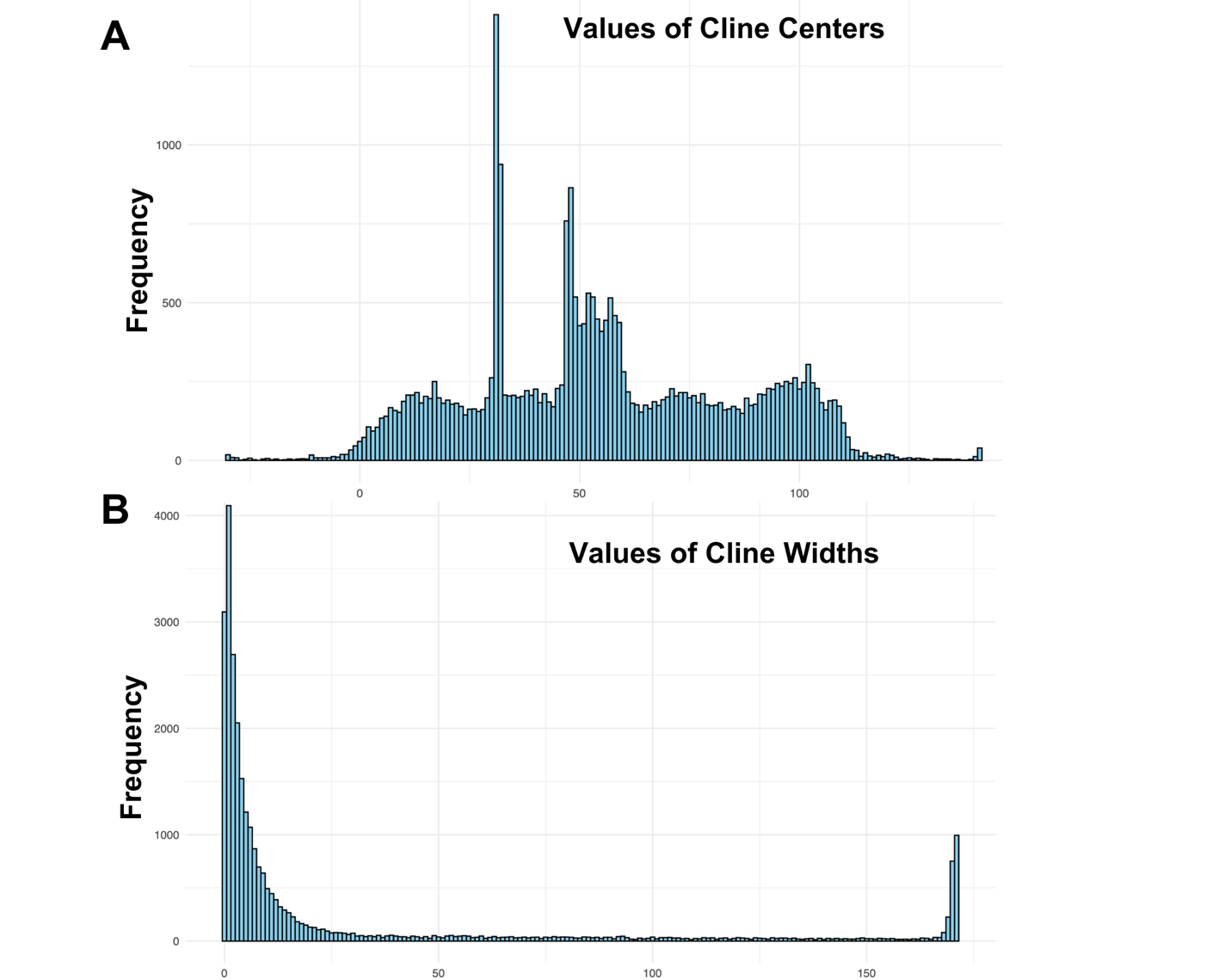

**Supplementary Figure S8:** Geographic cline properties. Highlighted in colors are the candidate found both with AMOVA and cline analyses (DFR and UDP-glycosyltransferase in pink and blue respectively) and in purple is the candidate found with GWAS analyses (7-deoxyloganetic acid glycosyltransferase).

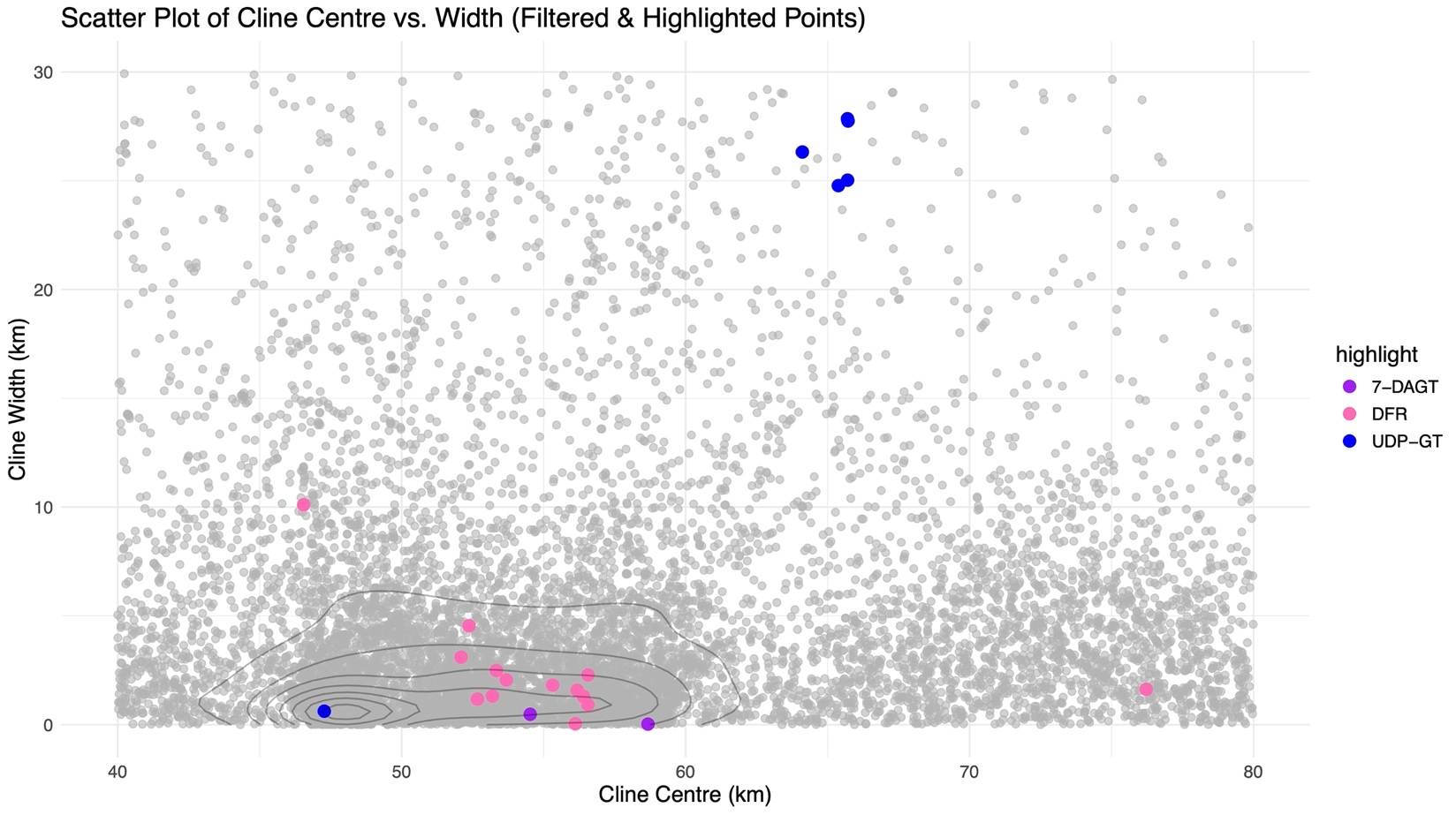

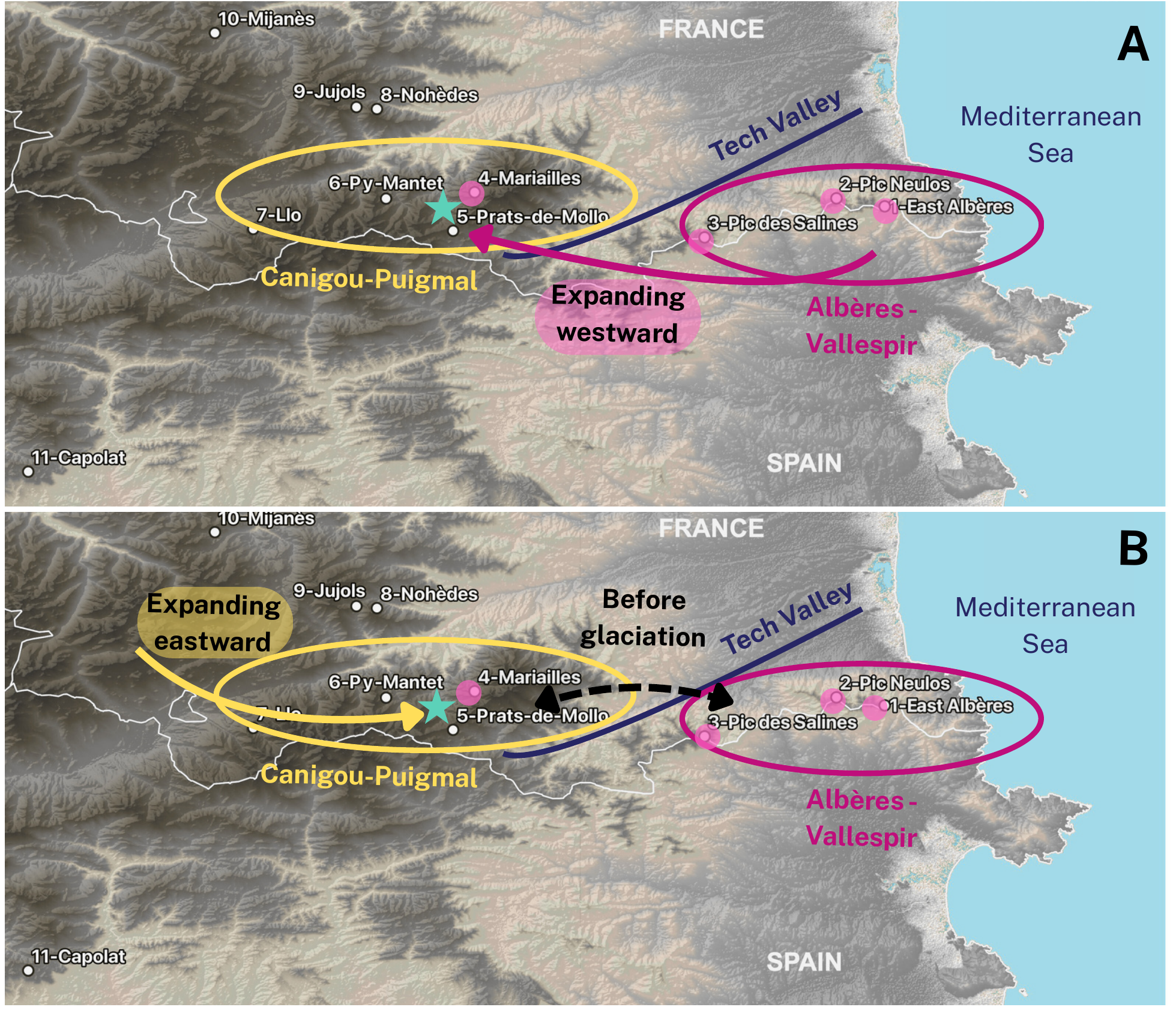
**Supplementary Figure S9:** Hypothetical scenarios for the origin and spread of the pink-flowered morph of *P. comosa* in the eastern Pyrenees. **A)** **Recent origin and westward expansion of the pink morph:** the pink-flowered morph originated recently in the Albères-Vallespir region (easternmost part of the range), possibly in response to new ecological conditions such as warmer, drier climates or higher UV exposure. This mutation (e.g., in FLS1 or regulatory elements upstream of DFR) might have been favoured by natural selection and began spreading westward toward the Canigou-Puigmal massif. The Tech Valley, a low-elevation and potentially unsuitable area for P. comosa, may act as a semi-permeable barrier to gene flow, slowing the westward expansion and contributing to the observed genetic structure. The current hybrid zone (green star) would thus result from a recent secondary contact between pink and yellow morphs. **B) Ancient divergence and secondary contact via eastward expansion of yellow morphs:** the pink morph was already present in the Albères-Vallespir and Canigou-Puigmal regions prior to the Last Glacial Maximum. These populations maintained gene flow across the region, including through the 4-Mariailles area. Following glaciation, gene flow between the Albères-Vallespir and Canigou-Puigmal may have ceased, possibly due to the Tech River acting as a barrier. Yellow-flowered populations, which may have persisted or recolonized from western refugia, then expanded eastward, leading to a secondary contact zone near Mariailles (green star). The resulting hybrid zone would reflect this older divergence and more complex demographic history. In both scenarios, the Tech Valley appears as a geographic and ecological boundary that may have shaped the formation and current position of the hybrid zone. Arrows indicate the direction of proposed population movements or gene flow.
